# Jointly representing long-range genetic similarity and spatially heterogeneous isolation-by-distance

**DOI:** 10.1101/2025.02.10.637386

**Authors:** Vivaswat Shastry, Marco Musiani, John Novembre

## Abstract

Isolation-by-distance patterns in genetic variation are a widespread feature of the geographic structure of genetic variation in many species, and many methods have been developed to illuminate such patterns in genetic data. However, long-range genetic similarities also exist, often as a result of rare or episodic long-range gene flow. Jointly characterizing patterns of isolation-by-distance and long-range genetic similarity in genetic data is an open data analysis challenge that, if resolved, could help produce more complete representations of the geographic structure of genetic data in any given species. Here, we present a computationally tractable method that identifies long-range genetic similarities in a background of spatially heterogeneous isolation-by-distance variation. The method uses a coalescent-based framework, and models long-range genetic similarity in terms of directional events with source fractions describing the fraction of ancestry at a location tracing back to a remote source. The method produces geographic maps annotated with inferred long-range edges, as well as maps of uncertainty in the geographic location of each source of long-range gene flow. We have implemented the method in a package called `FEEMSmix` (an extension to `FEEMS` from Marcus *et al*., 2021), and validated its implementation using simulations representative of typical data applications. We also apply this method to two empirical data sets. In a data set of over 4,000 humans (*Homo sapiens*) across Afro-Eurasia, we recover many known signals of long-distance dispersal from recent centuries. Similarly, in a data set of over 100 gray wolves (*Canis lupus*) across North America, we identify several previously unknown long-range connections, some of which were attributable to recording errors in sampling locations. Therefore, beyond identifying genuine long-range dispersals, our approach also serves as a useful tool for quality control in spatial genetic studies.

**Author Summary:** The movement of individuals across landscapes shapes genetic diversity and has significant implications for both evolutionary studies and conservation efforts. Advances in sequencing now allow researchers to analyze thousands of samples from broad geographic areas, helping to estimate local gene flow. However, long-range genetic flow can occur due to a host of reasons (e.g. natural weather patterns, migration for resources, etc.), and existing methods struggle to represent these patterns. In this study, we developed a method to identify and model these long-range genetic similarities as dispersals from a source to a destination over a landscape. In applying this method to over 4,000 human samples from Afro-Eurasia, we detected signatures of known long-distance dispersals from recent centuries. In applying this method to 100 gray wolf samples from North America, we found many unexpected long-range genetic connections, some of which turned out to be recording errors in sample locations. Thus, beyond detecting real long-range dispersal, our approach also serves as a useful tool for quality control in spatial genetic studies.

## Introduction

A key first step in understanding the genetics of a species is to understand its variation across the geographic range it inhabits (i.e., the geographic structure of genetic variation, or the “landscape genetics” of the species Storfer *et al*., 2010; Manel & Holderegger, 2013; Novembre & Peter, 2016; Bradburd & Ralph, 2019)). In many, or most species, isolation-by-distance patterns are common, in which genetic similarity is highest amongst the most geographically proximal individuals (i.e., Perez *et al*., 2018). The scaling of isolation-by-distance patterns often varies across a species’s range (“spatially heterogeneous isolation by distance”) due to factors such as persistent, prominent geographic features that alter migration, or non-equilibrium dynamics such as expansions from glacial refugia.

Models of spatially heterogeneous isolation-by-distance developed over the past decades have proven to be quite useful. In these models, local migration rates connecting neighboring local populations (or demes) are allowed to vary across the landscape. Methods based on such models can take geographic or ecological data as input and produce predictive maps of expected genetic differentiation (McRae, 2006; Dickson et al., 2019), or they can be used to fit regression weights for ecological factors that might affect observed genetic connectivity (Hanks & Hooten, 2013). The models can also be used with genetic data as input to produce maps of high and low levels of local gene flow (Petkova *et al*., 2016; Marcus et al., 2021), as is our focus here.

However, one limitation of models of spatially heterogeneous IBD is their inability to model long-range genetic similarity. In many taxa, localized gene flow is punctuated by pulses of long-distance dispersal across a landscape (Clobert, 2012). Such events produce cases where individuals separated by long geographic distances are remarkably genetically similar to one another. Ideally, methods to represent the geographic structure of genetic variation can also help identify when such long-range genetic similarities exist.

Being able to identify the location of the sources and destinations of recent putative long-range gene flow events can not only be useful in painting a fuller picture of the recent evolutionary history of dispersal and reproduction in a species, but can also be relevant for conservation in terms of identifying long-range genetic connectivity which may affect the long-term survival of populations (Rosenberg *et al*., 1997), and the functioning of ecosystems and the services ecosystems provide (Des Roches *et al*., 2018, 2021).

To address this challenge, we present an extension of the `EEMS` (“Estimation of Effective Migration Surfaces”) method from Petkova et al. (2016) and `FEEMS` (“Fast EEMS”) method from Marcus et al. (2021) that represent spatial genetic structure by inferring migration rates on the edges of a graph of connected nodes. The default graph is dense and has neighboring nodes connected to one another such that the resulting set of symmetric migration rates across the edges approximately specifies a continuous “migration surface”. The nodes in the graph represent local populations (i.e. sets of randomly mating individuals) and are referred to as demes for the remainder of the manuscript. On this graph, edges with low inferred migration help convey how samples are more dissimilar than expected given their geographic separation. Conversely, regions of high inferred migration depict when samples are more similar than expected given their geographic separation. We refer interested readers to Petkova et al. (2016) and Marcus et al. (2021) for a full explanation of the model (and to Lundgren & Ralph, 2019, for limitations of the model with regards to the modeling of directional/asymmetric migration).

Our extension, called `FEEMSmix`, detects when the nearest neighbor graph is insufficient to explain the data and adds directional long-range edges (LREs) to account for excess similarity between distant nodes in the graph. This similarity might arise from long-range gene flow or admixture events, or mistakes in record-keeping of the geographic position assigned to a sample. Long-range genetic similarity could arise due to a single recent, instantaneous pulse or from some form of continuous gene flow stretching into the past over a region of very low effective migration. In all cases, our method will represent these events in the form of an interpretable *source fraction* parameter, akin to an admixture proportion, from a hypothetical instantaneous long-range pulse that occurs just before sampling. This framework follows closely from existing methods that model the residual from an existing fit as a specialized admixture component (`TreeMix`, Pickrell & Pritchard (2012); `MixMapper`, Lipson et al. (2013); `SpaceMix`, Bradburd *et al*. (2016)). However, our method is novel in that it extends the `EEMS/FEEMS` framework, which models spatially heterogeneous isolation-by-distance. The new method models the observed long-range similarity in terms of a geographic source (i.e., a location with latitude and longitude), which aids in interpretation, while also having faster run times when compared to existing methods.

In this manuscript, we present the new method, `FEEMSmix`, and test it with simulations over a range of parameter values to quantify its performance. Finally, we apply this method to two large empirical data sets across different species and geographic ranges: first, to a data set of 111 wolf samples across North America (originally from Schweizer *et al*., 2016, and used as a prototype in `FEEMS`), and second, to a data set of 4,070 humans from over 300 sampling locations across Africa, Europe and Asia (compiled as part of Peter *et al*., 2020) to validate the working of the method in a well-studied system with findings that can be corroborated with alternative historical data sources.

## Results

### Analysis of a representative simulated dataset

We show a schematic workflow for the methodology of `FEEMSmix` using a representative simulation of a simple scenario of spatial population structure with a long-range gene flow event via Figure 1.

**Figure 1:**
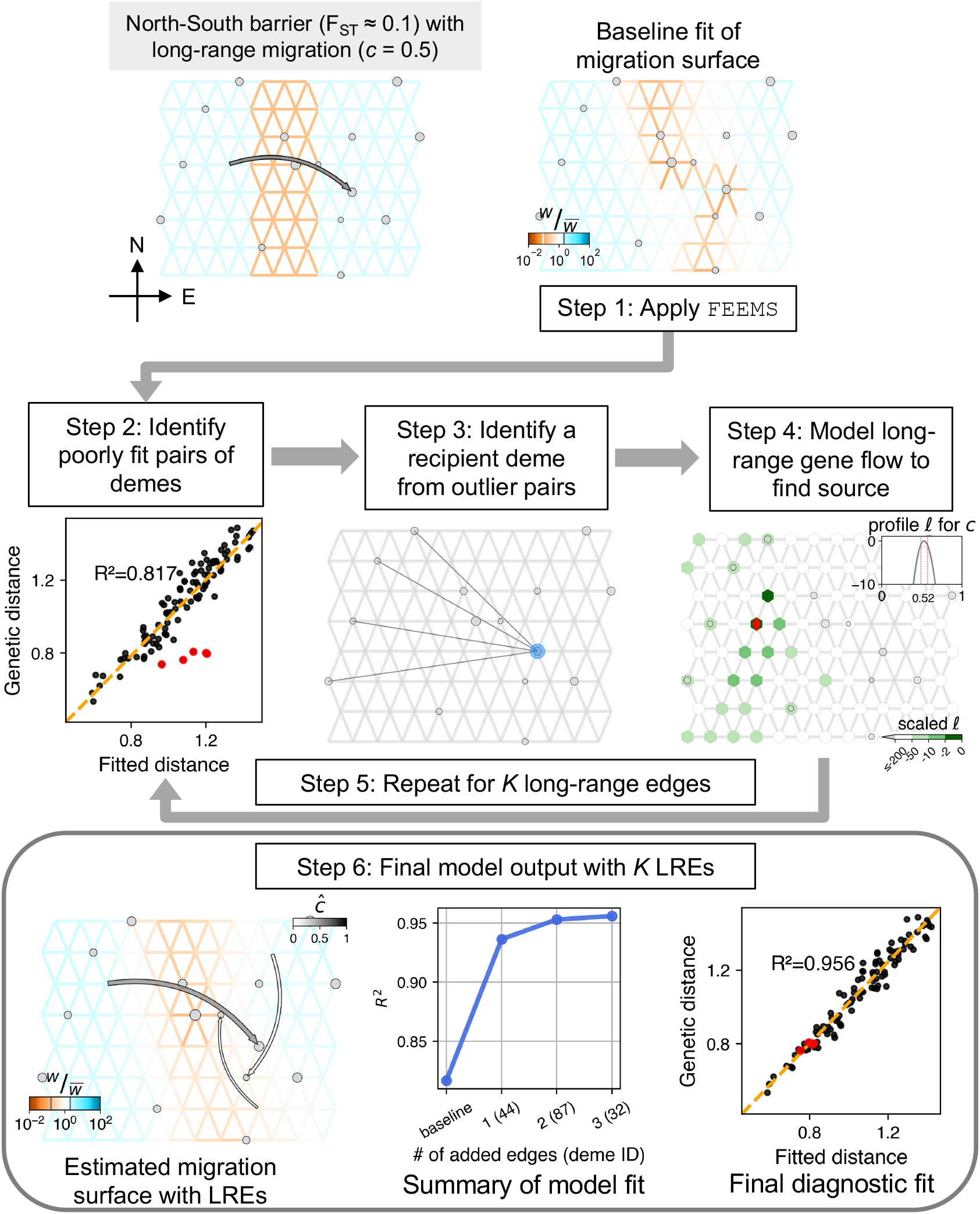
Summary of workflow of the `FEEMSmix` method presented via application to a representative simulation of a single barrier scenario with *variable, sparse sampling* (see main text for description of the steps and Figure S2 for the *constant, dense sampling* analog). In the simulated scenario (top left panel) and fitted results, the weights (*w*) on the edges that represent migration rates between pairs of demes are shown relative to their mean value across all edges (see legend for 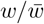).

For the simple scenario, we consider a spatial graph of locally connected demes that form a 8 *×* 12 grid with a barrier region of low migration running “North” to “South” across the center of the grid. The effective migration rates inside the barrier are ten times lower than the adjacent areas, resulting in an *F*_ST_ ≈ 0.1 across the barrier. The scenario includes an instantaneous long-range event going “West” (source) to “East” (destination) across this barrier just prior to sampling, with a backward migration fraction of *c* = 0.5. In other words, looking backward in time, 50% of the lineages in the destination deme migrate in a pulse-like fashion to the source deme. In `FEEMSmix`, we call this parameter the *source fraction* as it is always associated with a particular source and it represents the fraction of lineages that belong to this particular source going backward in time. To demonstrate the performance under two extremes of data availability, we use two sampling schemes called *constant, dense sampling* and *variable, sparse sampling* (shown in Figure S1). In the former scheme, we impose a 50% sparsity and sample the true source deme, with 10 individuals per sampled deme across the grid. In the latter scheme, we impose a strict sparse sampling to mimic real-world settings. More than 80% of the demes in the graph are unsampled, and we leave the true source deme of the long-range gene flow event also unsampled. In this case, we sample a random number of individuals between 1 and 10 per sampled deme. We note that in real data applications from continuous landscapes, users must choose the resolution of the grid to use for analysis, and helper functions in the software to help register their samples onto the grid. The full simulation parameters can be found under *Simulation* in Methods.

#### Step 1. Apply FEEMS

We fit the `FEEMS` method to the data using the deme-specific variance mode outlined in Marcus et al. (2021), with cross-validation to choose the appropriate tuning parameters *λ* and *λ*_*q*_. We refer to the resulting fit as the ‘baseline’ fit in the remainder of the paper.

In the example in Figure 1, we see that `FEEMS` does a mediocre job of reconstructing the low dispersal barrier region. In particular, it fits the regions poorly between the destination and source of the long-range gene flow event – as it compensates for the high genetic similarity between the source and destination demes on either side of the barrier by fitting a corridor of high gene flow between the two demes.

#### Step 2. Identify candidate poorly fit pairs of demes

In this step, given our motivation to detect and describe long-range genetic similarity that is not well represented by the `FEEMS` model, we identify pairs of demes that have a smaller observed (genetic) distance than expected under the `FEEMS` model. We calculate residuals between the observed and fitted genetic distances on a log scale. We designate the top 1% of pairs with the highest negative residuals as candidate outliers.

Even though we see a reasonably high *R*^2^ ≈ 0.82 in this representative example (Fig 1, Step 2), there are obvious pairs of demes that are not well fit under the model (indicated by the red circles, Step 2 panel).

#### Step 3. Identifying a putative recipient deme from outlier deme pairs

As a visual aid, we draw edges between the candidate outlier deme pairs identified in Step 2. As shown in this representative example, a single simulated long-range dispersal event between a source and destination deme will typically cause multiple demes neighboring the source to be fit poorly by the baseline `FEEMS` model. While this provides a summary of the `FEEMS` fit that emphasizes potential unmodelled long-range genetic similarity, it is useful to attempt to model a long-range gene flow event between a single single source deme to a single recipient deme. This is a compact way of representing and explaining the signal observed across multiple outlier pairs. To do so, we first need to identify putative recipient demes from the candidate outlier pairs.

We consider each outlier edge as specifying a potential long-range gene flow event and assess the favored directionality of gene flow for each outlier edge. We use the following algorithm: for a pair of sampled demes *i* and *j* implicated as an outlier pair, we fit them as the result of long-range gene flow (see Methods) in both possible directions, i.e., *i* → *j* or *j* → *i* with the → denoting the direction of gene flow from source to destination forward in time. If the model fit for *i* → *j* has a log-likelihood 2 units larger than *j* → *i*, we take *j* as a putative recipient deme and *i* if the opposite is true. In cases where these quantities are within 2 units of each other, both *i* and *j* are added as putative recipients to our list.

Taking this approach, the true destination deme is often found to be the putative recipient across multiple outlier pairs (Step 3 panel in Figure 1). The size of the blue circle shows the number of times this deme is picked as a putative recipient across all outlier pairs. Thus, we take the deme most often implicated as a recipient to be the first recipient deme to fit in Step 4, and in the case of a tie, the implicated deme with the most negative residual is chosen as the putative recipient.

#### Step 4. Fitting the source location for a chosen destination

For the chosen recipient deme from Step 3, we model a long-range gene flow event and find the maximum-likelihood estimate (MLE) of the source location and corresponding genetic ancestry proportion derived from that source (i.e., source fraction *ĉ* in Figure 1, Step 4 panel, also see Methods). We also output several visual summaries relevant to the fitting of the source location for a single LRE: *marginal likelihood surface* over the entire grid for the putative source location of the fitted LRE with darker green reflecting a higher log-likelihood of a particular deme being the source; *arrow from MLE source to destination deme* colored by *ĉ* (gray-scale from white to black for [0, 1]); *profile log-likelihood at MLE source* in gray for the estimated source fraction with dashed red lines indicating 2 log-likelihood units around the MLE and lighter grey lines in the background represent profile likelihoods for other potential sources that lie within this threshold.

For the example shown in Figure 1, the method’s first LRE identifies the true recipient deme and fits the source to be one deme away from the true source (shown as a red diamond) with a reasonably accurate estimate of the source fraction (*ĉ* = 0.52 versus *c* = 0.5 simulated).

#### Steps 5 - 6: Additional edges and final model output

After the fitting of the first LRE to the putative recipient deme, Steps 2 through 4 are repeated sequentially on a re-estimated migration surface that *contains* preceding LREs for a user-specified number of edges, *K* (see Methods for details).

The final output is a map that shows long-range arrows indicating source and destination for each LRE, colored by the associated MLE source fraction (*ĉ*), placed over the underlying grid with edges colored by their estimated parameters from a joint fit (Figure 1, Step 6, see Methods). The size of the arrow decreases with each added LRE in order to visually highlight the demes with the largest residuals. Akin to the practice for adding migration edges in `TreeMix` (Pickrell & Pritchard, 2012), we do not impose a strict stopping criterion, though users may find it helpful to inspect various outputs when interpreting their results and evaluating the number of edges to include in the final output. To aid in this inspection, the method outputs a plot of the *R*^2^ for fitting distances as a function of *K*. It is of interest to see whether adding edges continues to increase *R*^2^ substantially, and to note the smallest value of *K* for which there is a plateauing of *R*^2^. The method also produces a scatter plot of the observed genetic distance versus fitted distance *after* modeling outliers *and* fitting the background `FEEMS` parameters to show the improvement in fit via *R*^2^ (Figure 1, Step 6).

Also, because the long-range event accounts for the residual genetic similarity between the two sides of the barrier, the migration weight estimation on the graph improves, especially in the area between the source and destination demes. We also observe a higher *R*^2^ value compared to the baseline `FEEMS`, and an improved fit of the previous outlier pairs (red points, Figure 1).

We also show two additional LREs beyond the first edge. These LREs only marginally improve the model *R*^2^, with a plateauing after *K* = 1. In this example, the estimated source fractions for these LREs are low indicating weak signals (*ĉ* < 0.15), but they still represent genuine residuals from the baseline `FEEMS` fit. We observe a similar pattern across a suite of simulation replicates (see Figure S3).

### Evaluation across multiple replicate simulations

We apply the method to fifty replicates of the simulated scenario in Figure 1 and evaluate performance in terms of: **1)** the rate of finding the correct destination deme, **2)** the average distance from the MLE to the true source deme, and **3)** the error in the estimated source fraction (*ĉ*). We repeat this procedure across the two sampling schemes noted above, and for varying values of *c* (0, 0.05, 0.25, 0.5).

In the case of no long-range gene flow (*c* = 0), we find minimally biased estimates across the top LRE inferred by the method in both dense and sparse sampling scenarios (0.001 [0, 0.002] and 0.008 [0.003, 0.013] respectively, in Figure 2C).

**Figure 2:**
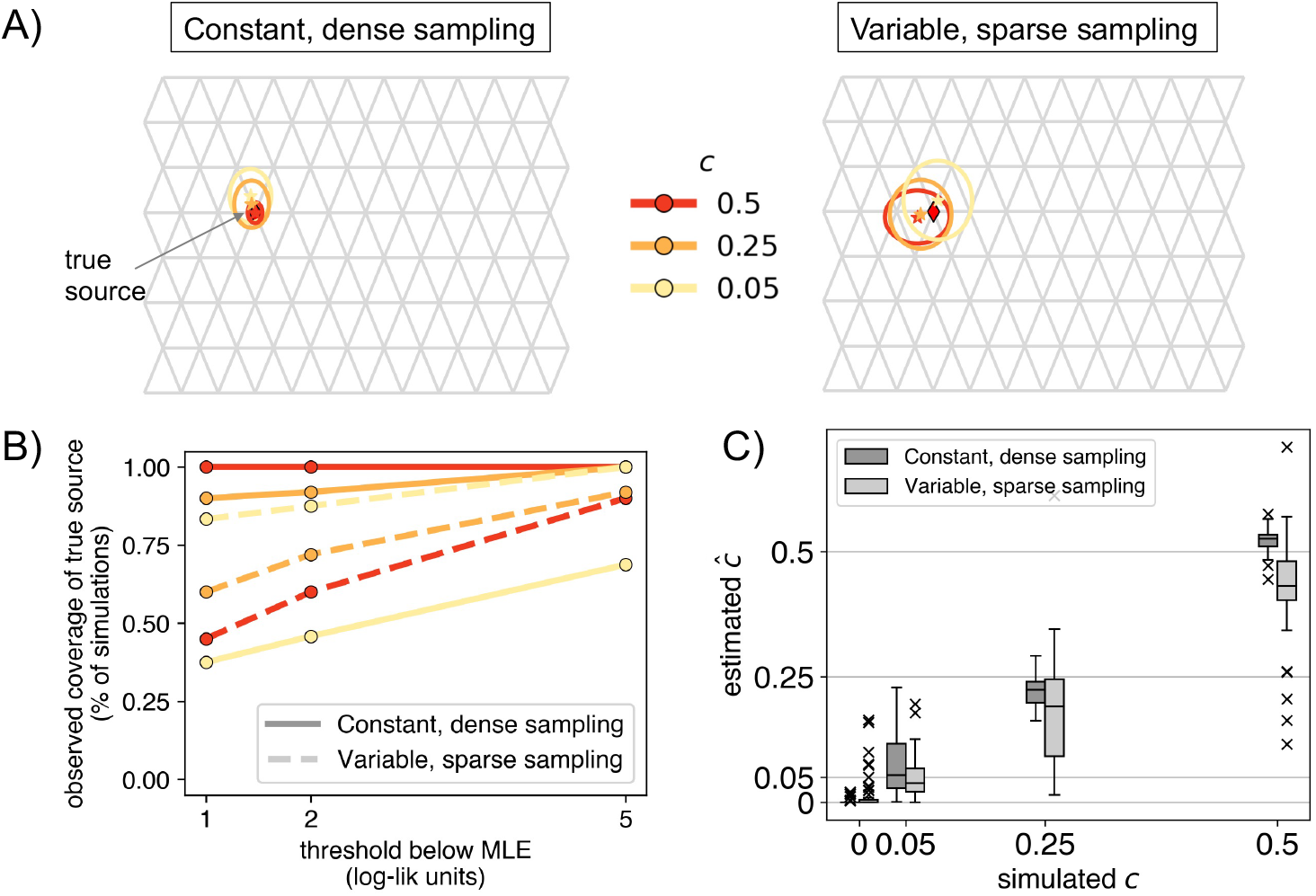
Range of performance metrics over 50 simulation replicates for each of the two sampling scenarios: **A)** Mean inferred location represented by a star with paired 2× standard errors on the mean location represented by boundaries of ellipsoids. The true source is indicated as a black diamond on each grid. In the sparse sampling scenario, we see that with stronger admixture (*c* = 0.5), the method tends to have smaller error when locating the source (since it is easier to locate a stronger signal), **B)** Coverage behavior of the log-likelihood surface, i.e., the percentage of simulations in which the true source was within a certain *x* threshold of the MLE source location, and **C)** Standard boxplots of the estimated MLE *ĉ* across all replicates for each simulated value of *c*.

Across both simulated scenarios, the method identified the true destination deme in 100% [93%, 100%] of the replicates with dense sampling and high gene flow (*c* = 0.5), and in 92% [81%, 98%] of replicates with sparse sampling. In the cases with low gene flow (*c* = 0.05), 0% [0%, 7%] of replicates had the destination recovered correctly in the dense sampling scenario, and 12% [5%, 24%] correctly in the sparse scenario. This seemingly better performance with sparser sampling is plausibly due to the fact that the sparse scenario has fewer sampled demes to choose from in the area surrounding the destination of the long-range event.

For evaluating the identification of the source, we fixed the destination deme to its true value and fit a source location (Step 4). When fitting cases with *c* = 0.5, the method was correct for 96% [86%, 100%] of the simulations in the dense scenario and 20% [10%, 34%] simulations in the sparse scenario. But, in the latter case, `FEEMSmix` picks a neighboring deme to the true source deme in 75% [62%, 87%] of the simulations (see Figure S3 for the suite of results). This accuracy is visually summarized in Figure 2A. As expected, uncertainty increases with weaker fractions and sparser sampling, though the MLE remain concentrated relative to the size of the grid, indicating that `FEEMSmix` is quite accurate at locating the true source. In Figure 2B, we assess accuracy using a coverage statistic, i.e., the percentage of simulations in which the true source is within a threshold *x* log-likelihood units of the MLE source. We see that for *c* = 0.5 in the dense scenario that this coverage is 100% [93%, 100%] for 2 log-likelihood units, and in the sparse scenario, it is 60% [45%, 74%]. The performance of the estimated source fractions is shown in Figure 2C. We estimate a small total bias in both scenarios (bias = 0.007 [0, 0.015] and −0.048 [−0.064, −0.033], for *c >* 0) with a higher bias in the sparse scenario.

### Application to North American gray wolves (*Canis lupus*)

We applied the method to the North American gray wolves data set of 111 individuals genotyped across 17,729 SNPs from Schweizer et al. (2016) used in the original `FEEMS` publication (Marcus *et al*., 2021). North American gray wolves are a highly mobile species (Schweizer *et al*., 2016) that show patterns of population structure consistent with some IBD (Musiani & Randi, 2024). Moreover, from a practical standpoint, this data set also provides a useful testing ground for our method given the sparsity of the sampling (that is typical for ecological studies conducted in the wild) and the prominent geographical features that introduce spatial heterogeneity in baseline effective migration rates across a broad continental scale. Further, dispersal patterns of these wolves are difficult to study due to the practical complexity in sampling such large areas while also accounting for external factors like seasonal, non-reproductive migrational patterns (Musiani *et al*., 2007; Mech, 2020).

The baseline `FEEMS` fit with the deme-specific variances and (*λ*_CV_, *λ*_*q*,CV_) = (2, 10) is shown in Figures 3A and S4 (the result is similar to the result in Marcus *et al*., 2021, which uses a single fixed variance). The model fits the regions encompassing geographical features like the the coastal mountain ranges in British Columbia, major waterways, and the tundra/boreal forest transition as having lower effective migration (model fit of *R*^2^ ≈ 0.95).

**Figure 3:**
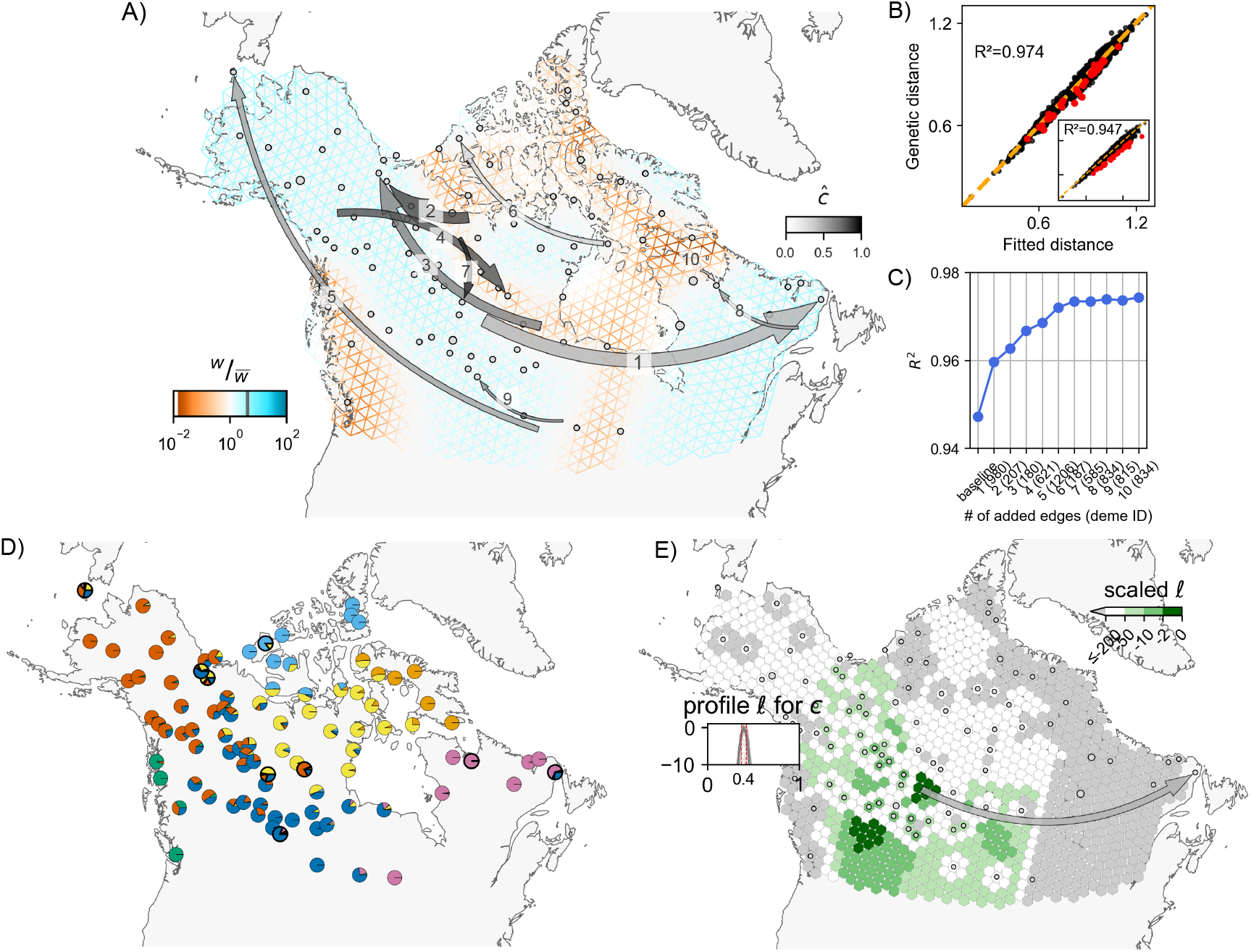
**A, B, C)** Full suite of `FEEMSmix` results with *K* = 10 for 111 North American gray wolves from Schweizer *et al*. (2016). **D)** The average of individual admixture proportions for each deme from an `ADMIXTURE` *K* = 7 run. The nine demes from subfigure **A** are outlined in black. **E)** Inferred surface for a particular long-range event to destination deme *980* (an example of an outlier that could be explained by admixture proportions) with *ĉ* ≈ 0.4 [0.37, 0.42] from `FEEMSmix`. The parameter *ĉ* reflects the estimated fraction of lineages from a particular source necessary to explain the observed long-range genetic similarity between deme *980* and the MLE source.

We ran the *iterative* fitting scheme over *K* = 10 edges, and found 9 unique recipient demes as a result. Each of these unique demes contained just a single individual. The identity of at least three of these recipient demes (*402, 621, 1206*) was not entirely surprising, as the original Schweizer et al. (2016) study had classified individuals belonging to these demes as being putatively admixed based on their ancestries from multiple sources in an `ADMIXTURE` analysis. In a similar vein, Figure 3D (also see Figure S5), shows that 8 of 9 demes identified by `FEEMSmix` (outlined in black) are modeled by `ADMIXTURE` with substantial proportions of ancestry from multiple sources, and each of these 8 demes appear distinctive from their surrounding demes in their ancestry profile. With close inspection, we find the LREs depicted by `FEEMSmix` generally help show pairs of distant populations in the `ADMIXTURE` plot that share ancestry in the same source populations.

To understand the inferred LREs in more detail, we examined sample meta-data. We assessed whether batch effects due to shared season of sampling (or year of sampling) could explain any of the long range similarities, but we found no such systematic associations (results not shown). However, we did find that 5 of 9 outlier samples have clear sampling issues or at least issues that raise concern; i.e., 2 of 9 were found to have recording or clerical errors in their locations, and 3 of 9 were recorded at vague or questionable locations (see Figure 4, and Section S1.3 in the Supplementary Material for further discussion).

**Figure 4:**
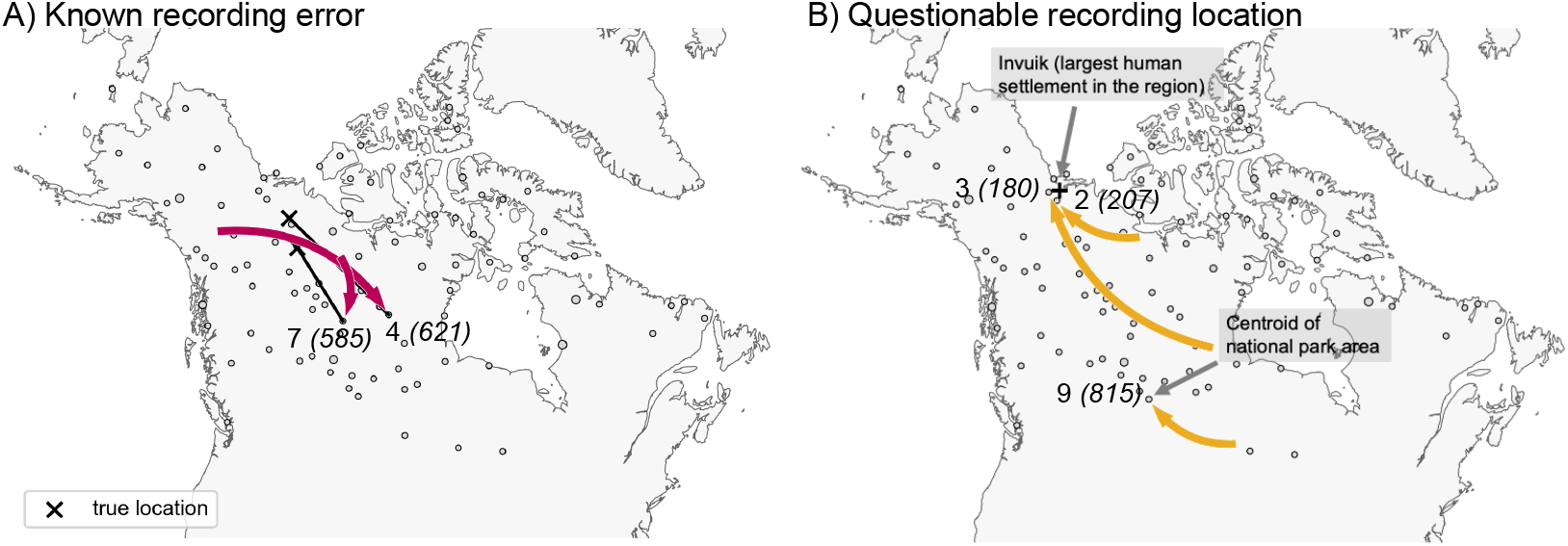
Results from examining the LREs found by `FEEMSmix` for potential explanatory factors. The figure shows several LREs labeled by edge ID and deme ID from Figure 3 and interpretations based on an investigation of the sample meta-data (see Section S1.3 in the Supplementary Material). **A)** For two samples, recording errors were discovered upon close inspection, and ‘X’ marks the corrected location of each sample. **B)** For three samples, the recorded locations are plausibly central locations where a sample was recorded as originating from, though the actual sampling event took place further away.

For comparison, we also ran two previously published methods to better contextualize our results: `SpaceMix` (spatial and with similar methodology for modeling admixture, Bradburd *et al*., 2016) and `TreeMix` (non-spatial but with very similar methodology for fitting residuals, Pickrell & Pritchard, 2012) (see Section S1 in the Supplementary Material for more details). We observed largely overlapping results amongst the three methods but found that `FEEMSmix` captured the largest set of unique long-range “dispersal” events (even with *K* = 10), whereas the other methods only found a subset of these putative events.

With regards to runtime on the wolf data, `FEEMSmix` took 2.5 hours for the entire workflow (with the baseline fit in `FEEMS` taking 15 minutes and fitting of the 10 LREs taking a total of 2.3 hours). In comparison, `TreeMix` took 32 hours to fit 15 edges and `SpaceMix` took 5 hours when estimating just admixture source locations and 6 hours when estimating both admixture source and geo-genetic locations (recommended usage). Both `TreeMix` and `FEEMSmix` will scale proportionally with the number of edges added to the baseline fit whereas the convergence in `SpaceMix` will depend on the complexity of the sampling posterior. We also note `FEEMSmix` can be run in parallel if each long-range edge is fit independent of other ones (i.e., in an alternative mode).

As one additional indication of the explanatory power of the `FEEMSmix` approach, for the wolf data, we carried out an experiment where we obscure the location of a particular individual from the model, and then re-assign this individual using the likelihood function of the model. We repeat this for each individual in the data set. This mimics spatial assignment of samples of unknown origin (e.g. museum specimens or retrieved biological material from smuggling, in the style of Wasser *et al*., 2004). Our results show that spatial assignment using the `FEEMSmix` likelihood function recovers sample positions similar to or slightly more accurately than a recently developed method (Battey *et al*., 2020) for the same task (see Figure S6).

### Application to humans (*Homo sapiens*) from across Afro-Eurasia

We also applied our framework to 4,070 modern humans from across 319 unique locations in Afro-Eurasia. This dataset was first compiled by Peter et al. (2020), and serves as a useful test bed for our method due to its broad geographic sampling, and more importantly, the ability to corroborate signatures of long-range genetic similarity with prior knowledge from both genetic and non-genetic sources, including archaeology, anthropology, and linguistics (Jobling *et al*., 2003)

First, we applied `FEEMS` to this data and found many previously-known patterns in human genetic variation, like elevated differentiation across the Saharan Desert, the Mediterranean Sea, and the Himalayan mountain range (Figure 5A for baseline `FEEMS` model with (*λ*_CV_, *λ*_*q*,CV_) = (3, 1)).

**Figure 5:**
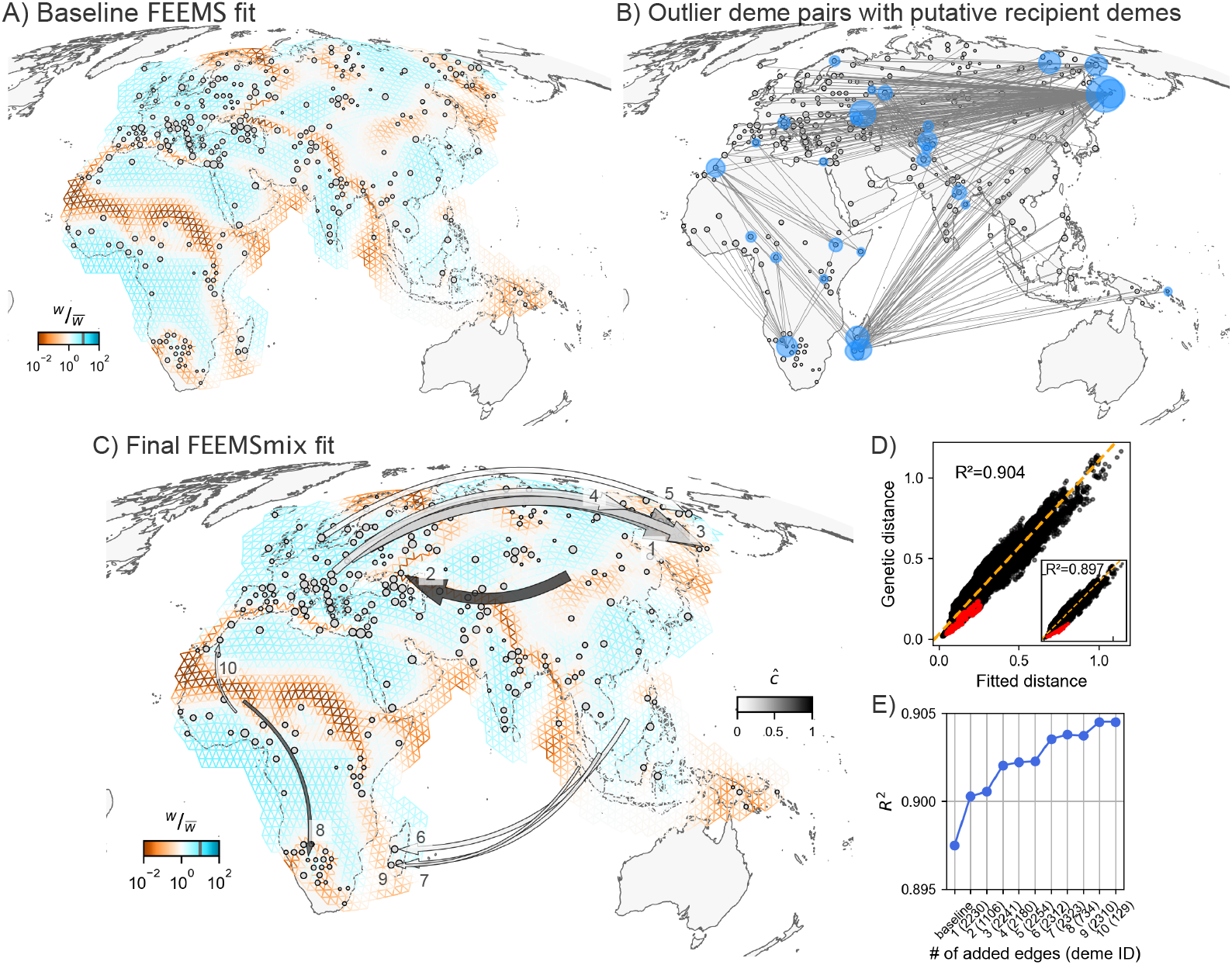
Empirical results from `FEEMSmix` for 4,070 humans from Peter et al. (2020). **A)** Baseline `FEEMS` fit for the 271 sampled demes with inferred migration troughs reflecting areas of historically low migration (e.g. Saharan Desert, Himalayan mountain range, Mediterranean Sea). **B)** A map showing the top 1% of outlier pairs, with putative recipient demes highlighted in blue by `FEEMSmix`. **C, D, E)** Full suite of `FEEMSmix` results with *K* = 10 LREs.

We run the same *iterative* fitting procedure as before for *K* = 10 LREs and find 10 unique recipient demes. All ten recipient demes contain individuals from a single population label in the sample meta-data.

The resulting ten long range edges fit by the method (shown in Figure 5C) can be interpreted in terms of five major signals, which we discuss in terms of the first added edge:

1. Aleut individuals from the Kamchatka peninsula in Russia (from Mallick *et al*., 2016) with a source fit from western Russia, with *ĉ* ≈ 0.5 [0.49, 0.52] (LRE 1). This long-range genetic similarity is supported by previous mtDNA results (Crawford *et al*., 2010) and `ADMIXTURE` results which show that about 20% of ancestry can be attributed to an ancestry found in northern Europe in models with *K* = 3 − 12 (from the original publication and Figure S7). We see similar signals for the Tlingit individuals from Mallick et al. (2016) (LRE 3), Yukagir individuals from Rasmussen et al. (2010); Yunusbayev et al. (2015) (LRE 4), and Chukchi individuals from Rasmussen et al. (2010); Cardona et al. (2014) (LRE 5) with decreasing strengths of northern European admixture (see Mallick *et al*., 2016; Sikora *et al*., 2019, and Figure S7);
2. Kalmyk individuals from eastern Russia (from Yunusbayev *et al*., 2015) with a source fit from the region of Mongolia and *ĉ* ≈ 0.8 [0.77, 0.82] (LRE 2). Kalmyks are a Mongolic-speaking group residing in Europe, with recent origins from East Asia with the latest wave of migration reported as happening in the 17th century (Nasidze *et al*., 2005);
3. Vezo individuals (north-western deme) from Madagascar (from Pierron *et al*., 2014) with source fit in southeast Asia with *ĉ* ≈ 0.12 [0.1, 0.13] (LRE 6). This LRE aligns with a well-known ancestral long-range dispersal across the Indian Ocean from south-east Asia with plenty of supporting genetic (Pierron *et al*., 2014, 2017) and linguistic (Adelaar, 2009; Beaujard, 2011) evidence. We see similar geographic signals with similar strengths replicated for the other Madagascar populations as well (Antemoro in the south-eastern deme (LRE 7) and Mikea in the south-western deme (LRE 9));
4. Bantu Herero individuals from Botswana or Namibia (from Mallick *et al*., 2016) show a source from western Africa with *ĉ* ≈ 0.73 [0.71, 0.74] (LRE 4). This signal is also backed by an `ADMIXTURE` analysis at *K* = 13 that shows a western Bantu component maximized in these individuals in Montinaro et al. (2017);
5. Moroccan individuals from south Morocco (from Henn *et al*., 2012) a source fit proximal to coastal Ghana and Nigeria with *ĉ* ≈ 0.23 [0.21, 0.25] (LRE 10). Henn et al. (2012) found that the southern Moroccan individuals are closest to the Luhyan population from Kenya (this relationship is also replicated in the initial outlier pairs found by `FEEMSmix` for this deme in Figure 5B). However, Henn et al. (2012) also infers that a migration to Morocco from sub-Saharan Africa occurred about 1,200 years ago, which coincides with the rise of the Ghanaian Empire that was involved in the trans-Saharan slave trade. This claim has gained further support more recently using haplotypic segments in Hellenthal *et al*. (2014) — and perhaps corroborates the placement of a source in western Nigeria from `FEEMSmix`.

Finally, the outlier deme pairs and the inferred source locations from `FEEMSmix` (in Figure 5B and C) are also supported by the geographic distribution of admixture *f*_3_ statistics (shown in Figure S8 for a few selected demes, computed using `AdmixTools` from Patterson *et al*., 2012).

## Discussion

In this paper, we present a method called `FEEMSmix` that represents the geographic structure of genetic variation using simultaneously a landscape of spatially heterogeneous gene flow and long-range gene flow events. It is built upon a previous method called `FEEMS` (Fast Estimation of Effective Migration Surfaces by Marcus *et al*., 2021), and follows in the same naming tradition of modeling residuals to baseline fits of the observed genetic data with a parameter specifying the strength of an instantaneous admixture pulse (e.g. `TreeMix`, Pickrell & Pritchard (2012); `MixMapper`, Lipson et al. (2013); `SpaceMix`, Bradburd et al. (2016)). However, we note that since we use the same approximate coalescent-time-based likelihood formulation as in `EEMS` (Petkova *et al*., 2016), this method could also rightly be called `EEMSmix`.

We have examined the sensitivity and accuracy of this method in simulations that mimic real-world settings, most importantly extremely sparse sampling. We also demonstrated the ability of the method to recover known signals of long-range gene flow events in a large empirical data set of humans across Afro-Eurasian panel. While we focused on *K* = 10, we do not think that these are the only instances of long-range dispersal in human history. For example, we find many instances of previously understood long-range genetic similarities for values of *K* between 10 and 25 (see Figure S9), e.g. Tiwari Brahmin from UP, India (LRE 12 shows a high proportion of Iranian ancestry, Nakatsuka *et al*., 2017), Hazara from Afghanistan (LRE 20 shows Central Asian ancestry, Mallick *et al*., 2016), and Masai from Kenya (LRE 21 shows Nilotic ancestry, Mallick *et al*., 2016).

In our analysis of 111 wolf samples across North America, we detect new signals of long-range genetic similarity, some of which appear to be genuine long-range dispersal, and others which appear to be artifacts of sampling. In many data sets one might expect small observational or recording errors to be on a scale that is negligible compared to patterns of IBD and inconsequential for most analyses. Here, the method helped identify two cases of mislabeled samples because they were identified as having unexpected long-range genetic similarities.

In empirical data, we find the method performs favorably to existing methods (`ADMIXTURE`, `TreeMix`, `SpaceMix`). Relative to these other methods, the framework developed here is unique in providing a single integrated framework for explicitly using geographic information to model varying rates of local migration as well as long-range admixture events. That said, running these other methods in a complementary fashion with `FEEMSmix` can help with corroborating results and gaining a more robust understanding of the genetic structure in one’s data.

As with any method that summarizes complex data using simple models and the notion of effective parameters, our method also comes with limitations that influence the interpretation of our results. First, our method `FEEMSmix` is built on top of `FEEMS` (Marcus *et al*., 2021), so it comes with the same underlying assumptions, which, in turn, influence the outliers that are chosen for analysis in `FEEMSmix`. Briefly, `FEEMS` requires a choice for the overlaid nearest-neighbor graph (“grid”) of assumed demes. It fits stationary, symmetric migration rates to the data, which can have drawbacks when asymmetric gene flow is pervasive (see Lundgren & Ralph, 2019).

Operationally, in terms of data, the method assumes the input are genotypes with no missing data and that all sampling locations are resolvable to the scale of the input grid of assumed demes. We also note that the choice of tuning parameters (*λ, λ*_*q*_) will affect the results from `FEEMSmix` in two ways: 1) the identity of potential LREs being implicated in the baseline `FEEMS` fit, and 2) the estimate of the resulting source fraction. As best practice, we recommend trying out different values of *λ* and *λ*_*q*_ spaced evenly on a log-scale around the baseline *λ*_CV_ value (e.g., {0.05 *× λ*_CV_, *λ*_CV_, 20 *× λ*_CV_}), when fitting the data. Typically, most outlier demes will persist through these different settings, but observing how the results change can be informative about the underlying signals in the data (see Figure S3 for replicates in simulations and Figures S4 and S10 for results with the empirical data sets). In simulations and in empirical data, the effect of changing *λ*_*q*_ is often negligible on both the outlier detection and the resulting source fraction estimates relative to the effect of the tuning parameter on the migration weights (see Figure S11 for a comparison of this in simulations).

In cases where the true history is not of an instantaneous gene flow event, the estimated source fraction *ĉ* should be viewed as an *effective* parameter that simply reflects the source fraction necessary to model the residual genetic similarity while accounting for a background of spatially heterogeneous local dispersal. In the case of an older pulse of gene flow, the expectation is an attenuation of signal over successive generations of background gene flow, causing *ĉ* to be downward biased (see Figure S12). Also, if there are multiple events from disparate sources to the same destination, `FEEMSmix` will typically choose the source with the most concentrated geographic signal though the log-likelihood surface could weakly reflect the presence of multiple sources depending on the background rates of gene flow (see Figure S13).

As an additional caveat, `FEEMSmix` only fits long-range gene flow events with destinations at sampled demes. If a gene flow event is truly recent and between unsampled demes, one will not be able to detect the instantaneous event in the data; however, if there has been sufficient time since the long-range gene flow event for the source lineages to traverse between unsampled and sampled demes, then `FEEMSmix` may potentially detect the event indirectly, depending on the background migration rates and time since event.

As a future extension to the method, it would be compelling to attempt a more precise modeling of older gene-flow events. This would require calculating the diffusion of lineages across the landscape in proportion to the estimated background equilibrium migration rates. Such a method could plausibly be applied to ancient DNA to calibrate migration surfaces in the past with only modern samples.

Another interesting extension would be to model gene flow as arising from a region rather than a single point source as is done here. One could imagine a model in which a set of sources contribute some lineages to a certain destination deme or set of demes. However, fitting such models would require searching over a large space of possible parameters, and it raises the issue of power in the data to discern among multiple plausible scenarios.

The ideal framework for understanding the genetic structure of any species would model time-varying dispersal regardless of its spatial scale with just a sparse sampling of individuals across the range. Though the method does not achieve this ideal goal, it takes a step in this direction, and in the meantime, provides a useful tool for describing the geographic structure of genetic variation by simultaneously illuminating long-range genetic similarity over a background of spatially heterogeneous patterns of isolation-by-distance.

## Methods

### Baseline FEEMS model

As the first step for our method, we fit the same model as `FEEMS` using the same input (genotype matrix and spatial locations for sampled individuals) and framework (assigning samples to closest nodes on a user-defined triangular grid, and estimating graph-specific parameters via penalized maximum-likelihood) with the same modeling assumptions (exchangability of individuals within a deme, symmetric migration, unlinked SNPs and multivariate normal assumption for the allele frequencies). We employ a modified version of `FEEMS` that allows for modeling deme-specific variance parameters compared to the default version that uses a single fixed variance across the grid. Although this version, originally presented in Marcus et al. (2021) but passed over for the faster, more parsimonious default version, increases the number of parameters by *o* (number of observed demes) and lengthens runtime, we found the added robustness when fitting long-range gene flow events to be a worthwhile trade-off. More specifically, this entailed replacing the fixed term, *σ*^2^diag (**n**^−1^), by a vector, **q** ∈ ℝ^*o*^, in the `FEEMS` likelihood (Equation 7 in Marcus *et al*., 2021) to parameterize deme-specific variances. Also, to avoid over-fitting, a penalty term is added to the likelihood, with scalar *λ*_*q*_ such that larger values impose greater similarity across the elements of **q** (see Marcus *et al*., 2021, for a complete description).

### Modeling long-range genetic similarity as instantaneous gene flow events

We model gene flow along each LRE as a uni-directional instantaneous pulse event such that a fraction *c* ∈ [0, 1] of genetic lineages in destination deme *d* descend from a source deme *s*. Within this model, the expected pairwise coalescent time *post*-event 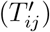 between any two demes *i* and *j* can be derived as a function of the expected pairwise coalescent times *pre*-event (*T*_*ij*_) and this source fraction *c*:

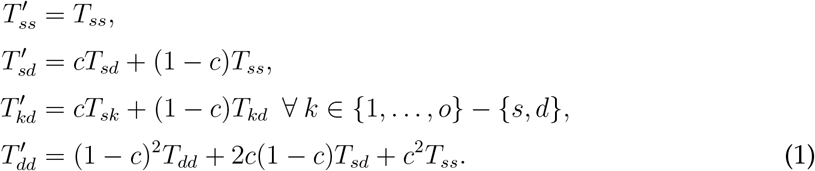

### Approximation of pairwise coalescent times in an unsampled deme

To model the expected pairwise coalescent times *post*-event in Equation (1) when the source is an un-sampled node (say, 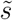) in the framework, we need an expression for the expected coalescent time within this deme, 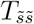, but the baseline `FEEMS` model only estimates this deme-specific variance parameter (which is related to the coalescent time, see Equation (3)) for sampled demes on the grid. As a result, we provide an approximation for the coalescent time *T* within an unsampled deme by spatially interpolating between values at sampled demes on the grid using a kriging approach with an exponential variogram model defined by *q*(*R*) = *b* + *C*_0_ (1 − exp(^−*R*^*/*_*a*_)), where *b* is the nugget, *C*_0_ is the sill and *a* is the range parameter. Notably, here, we use the resistance distances *R* instead of geographic distances to account for the effect of spatially heterogeneous IBD. These three kriging parameters are estimated by fitting the variogram to inferred variances from sampled demes across the grid by minimizing the squared difference between the expected and inferred values. The kriging weights *γ*_1,···, *o*_ for each of the observed demes are determined by solving the set of linear equations given by 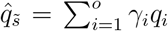 using weighted least squares optimization for a specific unsampled deme, 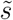.

### Specifying the FEEMSmix likelihood

As a piece of software, `FEEMSmix` is built completely on top of the `FEEMS` model developed by Marcus *et al*. (2021), but we use a parametrization of the model in terms of pairwise coalescent times developed in `EEMS` by Petkova et al. (2016). This is because each approach has unique advantages: 1) the `FEEMS` framework provides fast gradient-based optimization machinery for penalized likelihood-based optimization (whereas `EEMS` uses Markov chain Monte Carlo in a Bayesian framework), and 2) the `EEMS` likelihood parameterizes the genetic distance between samples in terms of pairwise coalescent times, which are more readily adapted for extending the model to more complex scenarios of gene flow.

To achieve this, we connect the `FEEMS` likelihood to the equivalent `EEMS` likelihood (Equation 3 in Petkova *et al*., 2016, or Equation (5) in this text), and restate the former in terms of expected pairwise coalescent times between lineages sampled from pairs of demes. First, we follow `EEMS` and define the expected symmetric unscaled genetic distance matrix **Δ** ∈ ℝ^*o×o*^ as a function of coalescent times that are approximated using a resistance distance matrix **R** and node-specific variance parameters **q** in the following way (for explanations of the approximations see the Supplement of Petkova *et al*., 2016):

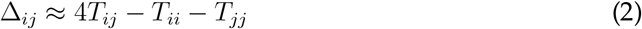

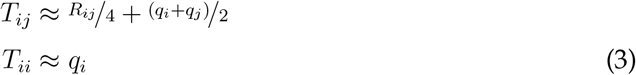

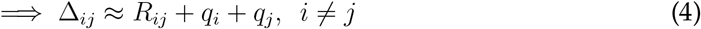

Importantly, the resistance distance between two demes is given by 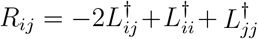, wherein **L**^*†*^ is the pseudo-inverse of the graph Laplacian given by L = diag (W1)−W (McRae, 2006; Hanks & Hooten, 2013; Marcus et al., 2021). In this way, we can see how the edge weights W and node variances q estimated in `FEEMS` are connected to the quantities in the expected genetic distance matrix in `EEMS`.

Based on the derivations above for the coalescent times after a long-range gene flow event, and applying the EEMS approximations that approximate coalescent times by a combination of pairwise commute times (i.e., resistance distances) and within-deme coalescent time parameters (*q*, Equations (1)-(4)), we can write equations for the elements of the expected genetic distance matrix *post*-event **Δ**′ as a function of *c*,

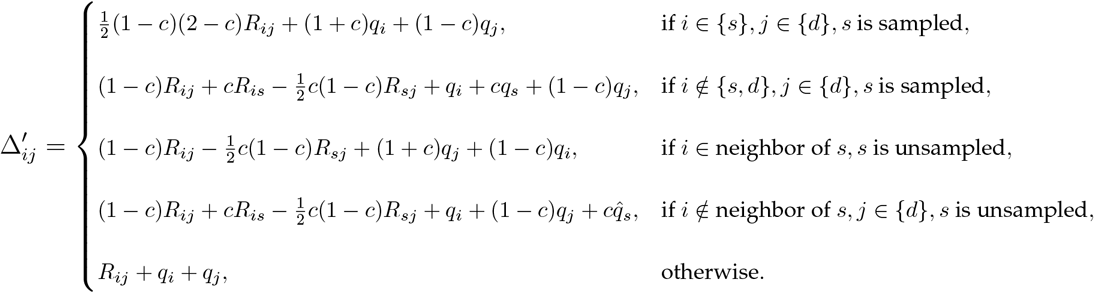

Note that for *c* = 0, these equations reduce to the baseline model given in Equation (4). We also use a separate equation (resembling Equation (1)) for the case when an unsampled deme is a nearest neighbor to a sampled deme, so as to ensure a smoother likelihood surface around sampled demes. When fitting *c*, we do not apply any penalty for sparsity (i.e., *ĉ* is the maximum value of the marginal likelihood for *c*).

With the expression for the expected pairwise distances (**Δ**′) in hand, we then follow `EEMS` and model the *observed* pairwise genetic distance matrix 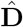 as a draw from a Wishart distribution centered on **Δ**′:

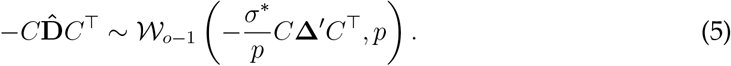

The matrix *C* is a contrast matrix to remove the overall mean. The *σ*^⋆^ is a scaling parameter, which we set to one, as we found its fitted value to be approximately equal to 1 across a range of simulation parameters, and forcing it to be 1 should just scale the units of **R** and **q** to be in units of expected distance. Finally, *p* is the degrees of freedom which is set to the number of unlinked SNPs as in Petkova et al. (2016). Consequently, we strongly recommend using only unlinked SNPs, as including linked SNPs may result in overly confident likelihood estimates. This issue is particularly salient in `FEEMSmix`, where the likelihoods are used to identify putative source demes across the grid.

### Joint optimization of the likelihood with a single long-range edge

Here, we outline the two-step procedure followed to fit a single LRE during Step 4 in Results. In our first ‘pre-fitting’ step, we fit a model of instantaneous gene flow from *every* deme in the grid to the putative recipient deme in an independent fashion. This is done by minimizing the negative log-likelihood of the data in Equation (5) for the source fraction *c* ∈ [0, 1], *while* holding the other parameters constant at their baseline values. This first step is very fast as for each source, we only optimize over a single dimension.For our second ‘re-fitting’ step, we choose only the top fraction (e.g. 1%) of demes with the highest log-likelihoods from the previous step to perform the joint fitting procedure in which we estimate *all* parameters in the model. This is an attractive approach as it saves us the effort of not having to estimate joint fits for demes that have a very low likelihood of being the true source deme (i.e., a vast majority of the demes in the grid), but still provides us a way of searching over the entire grid.

We employ a coordinate-descent approach when minimizing the negative log-likelihood of the model with *all* parameters for a particular LRE: edge weights, node variances *and* source fraction. Similar to the ‘pre-fitting’ step, we first optimize over the single dimension of the source fraction *c* using Equation (5) but with the parameters held constant at their values estimated from the baseline fit in `FEEMS`. Then, holding *c* constant at its MLE value, we re-fit the edge weights and deme-specific variances using the likelihood from `FEEMS`. Then, we repeat the optimization procedure for the source fraction holding the other parameters constant at their MLE values, and so on, until an absolute tolerance is reached in the values of the parameters (10^−3^ for the source fraction and 10^−7^ for the edge weights and deme-specific variances). We initialize the parameters at the baseline `FEEMS` fit to speed up convergence as we only expect slight deviations in estimates for any single LRE.

This two-step approach provides us with a fast and simple way to optimizing the joint likelihood as it allows us to reuse the flexible machinery formulated in Marcus *et al*. (2021) for computing the gradients as a function of the edge weights and deme-specific variances. The only substitution we perform is to replace the expected covariance matrix **Σ** in Equation 18 of Marcus et al. (2021) with our reformulated expected covariance matrix **Σ**′ that is simply derived from the expected distance matrix **Δ**′, 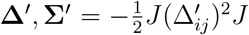 (where 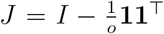 is the centering matrix and *I* is the identity matrix). Optimization in `FEEMSmix` is done using the Nelder-Mead and L-BFGS-B algorithms implemented in `scipy` (Byrd *et al*., 1995; Gao & Han, 2012; Virtanen et al., 2020).

### Joint optimization with multiple edges

By default, `FEEMSmix` fits multiple edges using an *iterative* approach. First, we fit a long-range edge to the putative recipient deme with the largest number of outlier pairs from the baseline fit using the workflow in Section, and then, we fit a LRE to the next putative recipient deme over a surface *containing* the previous LRE, and so on. We repeat this procedure until a user-specified *K* edges are fit on the baseline graph. As a precautionary measure against overfitting, we default to only allow for a maximum of two LREs to be fit to the same recipient deme. However, this can be changed in the software version of the package with a simple flag. We also do not specify a default stopping criterion here just as in `TreeMix`, as it is difficult to have rigorous, stable criteria, and we recommend users apply the method in a more exploratory manner.

Finally, we also provide users with the option of fitting multiple edges using an *independent* approach for each putative deme in the initial list of outliers from the baseline `FEEMS` fit.

### Simulation

All simulations were conducted in `msprime` (Kelleher *et al*., 2016). As a set of test cases, we simulate a 8 *×* 12 nearest neighbor graph/grid of demes with a barrier at the center of the grid to capture a spatially heterogeneous migration landscape. The longrange gene flow event occurs as an instantaneous pulse (`MassMigration` event) with varying source fraction *c* ∈ {0.05, 0.25, 0.5} across the barrier. We use 1,000 independent SNPs with both of the following scenarios.

### Constant, dense sampling

We sample 48 demes in the grid (50% sparsity, with the true source deme being sampled) with 10 individuals per deme and a uniform population size of 1,000.

### Variable, sparse sampling

Here, we randomly sample 15 demes across the grid leaving the true source deme unsampled, and with a variable number of individuals per deme drawn uniformly from between 1 and 10. We also simulated unequal population sizes across the grid, drawing from a uniform distribution between 100 and 10,000.

### Application to North American gray wolves (*Canis lupus*)

The data set of 111 individuals sampled across North America was originally collected in Schweizer et al. (2016) and used as an example data set in `FEEMS`. The SNPs were pruned for a 5% minor allele frequency cutoff and a 10% missingness rate, resulting in a total of 17,729 SNPs. Since this method is built on top of `FEEMS`, we used the same dense grid chosen in the original analysis to fit the baseline model. This grid had a cell area of approx. 6,200 sq. km. and a cell spacing of approx. 110 km. As a result of this grid choice, we see that the 111 individuals get assigned to 94 demes across the species’ continental range. Both `TreeMix` and `SpaceMix` were run on this sampled deme level so as to ensure a like-to-like comparison with the results from `FEEMSmix`.

### Application to Afro-Eurasian panel of humans (*Homo sapiens*)

We start with the data set of 4,302 individuals curated in Peter et al. (2020) for our analysis. From this data, we subset to those with public sharing permissions and are left with 4,070 individuals. This final data set consists of 19,954 SNPs from 319 distinct sampling locations across Afro-Eurasia. For this analysis, we use a grid with cell area of approx. 25,000 sq. km. and a cell spacing of approx. 220 km., resulting in a high sampling resolution with a total of 290 sampled demes across the grid.

## Data Accessibility

The wolves data set is provided as part of the `FEEMS` package in Marcus et al. (2021) (and is also publicly available from the original publication, Schweizer *et al*., 2016). The human data set was first compiled as part of Peter et al. (2020). The corrected wolves data set and the human data set used in this study can be found at https://doi.org/10.5061/dryad.p8cz8wb18. All simulated data can be reproduced using code in https://github.com/VivaswatS/feems/tree/admixture_edge. Finally, `FEEMSmix` is readily available as a complete `python` package from https://github.com/VivaswatS/ `feems`, and will be integrated into the `FEEMS` package upon publication.

## Author Contributions

VS and JN formulated the model. VS ran the simulations and analyses. VS, MM and JN interpreted the results and wrote the manuscript.

## Acknowledgments

We would like to thank members of the Berg, Novembre, and Steinrü cken labs, as well as members of the University of Chicago Program in Computational Biology (PCB) community for helpful discussions and feedback during the development of this project. We would also like to thank Rena Schweizer for help with a preliminary interpretation of the results. We would also like to thank Gideon Bradburd for comments on an early draft of the manuscript. Additionally, we thank Jeremy Berg, Matthias Steinrü cken, and Xuanyao Liu for support at all stages of this work. Computing was performed on servers maintained by the University of Chicago Research Computing Center. Funding to JN and VS was provided by NIH grants R35 GM149521 and R01 GM132383. MM was supported by European Union - NextGenerationEU, under the National Recovery and Resilience Plan (NRRP), Project title “National Biodiversity Future Center - NBFC” (project code CN 00000033).

## Notes

### Competing Interest Statement

The authors have declared no competing interest.

### Summary of Updates

The manuscript files have been updated to provide the supplemental information text.

