## Supplemental Information for "Jointly representing long-range genetic similarity and spatially heterogeneous isolation-by-distance"

### Supplemental Material for "Jointly representing long-range genetic similarity and spatially heterogeneous isolation-by-distance"

Vivaswat Shastry[1], Marco Musiani[2] and John Novembre[3]

[1] Committee on Genetics, Genomics and Systems Biology, University of Chicago, Chicago, IL, USA

[2] Department of Biological, Geological, and Environmental Sciences, University of Bologna, Bologna, Italy

[3] Department of Human Genetics, University of Chicago, Chicago, IL, USA

Corresponding author: John Novembre
920 E. 58th St.
Chicago, IL 60637, USA


Running title: Representing long-range genetic similarity

### S1 Results from North American gray wolf samples of Schweizer et al. (2016)

#### S1.1 Using SpaceMix

SpaceMix (Bradburd et al. 2016) was run in two modes: "estimating geogenetic and admixture source locations" and "estimating admixture source locations". As basic preprocessing and to ensure a like-to-like comparison, we clumped individuals into demes specified by the grid used by FEEMS and passed as input to SpaceMix the sample allele frequency matrix and the number of samples per deme.

In the first mode, the chain was initially burned in for 100,000 steps and then run for a total of 5,000,000 steps with a sampling every 2,000 steps to decrease autocorrelation. This produced 2,500 samples from the posterior distribution. . Under this scenario, SpaceMix does quite well to capture the broad geographic patterns in the data (similar to PCA in Figure S14B), but only implicates a single deme (*737*) with any significant amount of admixture ($>1\%$, Figure S17). A probable reason for this is that the residuals from other putative admixed sources are modeled away by their 'geogenetic' locations, so a long-range admixture event does not need to be evoked to fit the data (for example, see the locations of demes *621* and *1206*). However, the placement of the other putatively admixed demes (found by both ADMIXTURE and FEEMSmix) followed their locations in principal components space quite closely.

In the second mode, the chain was initially burned in for 500,000 steps and then run for a total of 35,000,000 steps with a sampling every 10,000 steps. This produced 3,500 samples from the posterior distribution. This mode is similar in idea to FEEMSmix, and we found largely overlapping patterns between the two methods (see Figure S18). For instance, six demes with stronger signals of admixture (median admixture proportion $>5\%$) were implicated in long-range events, and five were in common with FEEMSmix: *980, 1206, 621, 180, 207*. Here, the likely location of the sources were very similar in range to locations estimated by FEEMSmix, with the median admixture proportions being much lower than the MLE $\hat{c}$ from FEEMSmix, with varying degrees of similarity to the values obtained from the previous ADMIXTURE analysis.

Finally, convergence for both modes was verified by observing trace plots, acceptance rates and the joint marginal distribution of samples across the different stages of the MCMC. Both modes produced excellent fits to the observed sample covariances with $R^2 > 0.9$ (see Figure S16), indicating that SpaceMix does indeed explain the underlying spatial genetic variation in the data.

#### S1.2 Using TreeMix

TreeMix (Pickrell and Pritchard 2012) was run on the allele count matrix for the same set of SNPs across the 94 sampled demes for $m = 1$ to $m = 20$ migration edges using default parameters. We found no significant increase in the log-likelihood of the model on adding edges beyond $m = 15$, so we use the resulting topology with 15 migration edges for our interpretations in the main text (see Figure S15A).

In general, TreeMix does well to capture the broad geographic and ecotypic patterns present in the wolves (see Figure S15A). A few notable exceptions in the tree structure were the placement of deme *621* next to the WestForest populations and the location of deme *1206* as being an outgroup to the rest of the populations. With regard to the migration edges, 14 our of 15 edges were between Arctic and HighArctic populations and did not replicate patterns found by SpaceMix or FEEMSmix. But, interestingly, FEEMSmix does produce a similar pattern when run under a baseline FEEMS fit with a *single, fixed variance* across all demes (see Figure S4). These Arctic and HighArctic populations also showed the highest amounts of drift under the TreeMix model and since the deme-specific variance parameter in FEEMS captures a quantity proportional to the effective population size, we believe that by estimating this quantity, we model away any such residuals that TreeMix picks up (hence, just a single outlier from this region with under the default model of a *deme-specific variance*). The singular edge in TreeMix that is *not* between Arctic and HighArctic populations is actually from the base of several different populations to the branch leading down to the AtlanticForest populations of which *980* (largest outlier in FEEMSmix) is a part of. In short, TreeMix only seems to model this largest signal of long-range gene flow, but misses on the more subtle signals captured by ADMIXTURE, FEEMSmix and SpaceMix.

#### S1.3 Using FEEMSmix

We identified $K = 10$ long-range edges (LREs) upon *iterative* fitting of the wolves data with 9 unique recipient demes (see Figure 3A). To better understand the inclusion of these edges in the results, we investigated the sample meta-data. There, we found that 2 out of the 9 recipient demes contained samples with recording errors, and 3 out of the 9 recipient demes contained samples that were registered to questionable locations. During our investigation, we also found evidence of one more sample (*189*) as being misrecorded, and this was implicated as the $24^{th}$ LRE in our analysis (see Figure S20). We find that the inferred locations are in the same general direction as the true locations (though, in some cases, off by a few hundred kilometers, see Figure 4A). Deme *621* is implicated in a SpaceMix analysis that places its inferred source in the same direction as FEEMSmix, but much farther west and north than its true location (see Figure S18). Deme *189* is implicated as an outlier in TreeMix as well.

For the samples that were reported as coming from questionable locations, we found that sample *815* was assigned to the centroid of a national park area (indicating a proxy for the region for where it was found). With this sample, we found that the inferred locations in FEEMSmix are proximal to their recorded locations, which is a loose indicator that these wolves may habit this general area but may not actually be from this specific location (and, as a result, are found to be outliers by the method). For the other two samples (*180, 207*), we found in the sample meta-data that both these samples were registered in the town of Invuik (black plus sign in Figure 4B) in the Northwest Territories of Canada. This is the only town in the region, so we infer that these wolves were most likely hunted somewhere else and brought to this town for processing (and, eventually, registered). FEEMSmix infers the source locations for this sample to be southeast of this town. This is in the same general direction as the source inferred by `SpaceMix` (see Figure S18). A supporting result is that in `TreeMix`'s inferred tree (which is fit without any spatial information) the samples closest to the `FEEMSmix` MLE source deme and those from the destination deme are inferred to be sister populations.

We note that `SpaceMix` also implicates the 2 out of 3 samples that were identified as being recorded at questionable locations (demes *180, 207*) and 1 out of 3 samples that were results of recording errors (deme *621*), whereas `TreeMix` only finds one 1 out of 3 samples that were results of recording errors (deme *189*).

Out of the remaining 4 samples, 3 samples (demes *980, 1206, 187*) show admixed ancestry in an `ADMIXTURE` analysis (Figure 4B), of which 2 out of 3 (demes *980, 1206*) are also implicated in a `SpaceMix` analysis (Figure S17). Out of these two outliers, deme *980* is found to be the source of a long-range migration event in `TreeMix`, though deme *1206* is not. Based on the inference of partial admixture memberships to multiple ancestries to deme *1206* and its basal outgroup position in the tree (and its location as an isolated sample on the westernmost island off the coast of Alaska), we hypothesize that this sample is likely a descendant of an ancestral wolf population that spread eastwards and gave rise to the populations that were eventually sampled from the central and eastern parts of the continental range. (This represents a model mis-specification under both `FEEMSmix` and `SpaceMix`.) Finally, the one deme that is a somewhat puzzling inclusion in `FEEMSmix` is deme *834* which doesn't show multiple memberships in an `ADMIXTURE` analysis, but still requires two LREs with different sources to model away the remaining residual under the model.

Finally, we present a reanalysis of the wolf samples with the locations corrected and the ambiguous samples removed in the section below.

##### S1.3.1 Re-analysis of the corrected wolf samples with `FEEMSmix`

After the correction, we are now left with 108 wolf samples spread over 89 unique nodes in the graph, where $\sim 90\%$ of nodes contained just a single sample. The estimated migration surface over these samples (Figure S20A) is very similar to the migration surface estimated in the main text. This baseline fit also provides an excellent fit to the corrected data $R^2 \approx 0.96$.

Out of these 9 demes, we found that 3 belong to the outliers from our original analysis in the main text. For two of these demes, the inferred source location and fraction are found to very similar to the estimates from the original analysis. However, for deme *834*, the inferred source area is found to be quite west of its original location, though the inferred source fraction is quite low for this LRE ($\approx 0.1$ $[0.08, 0.13]$).

Out of the remaining 6 demes, we found that 2 demes contained reassigned samples from our original analysis (demes *336* and *213* were reassigned demes *585* and *189* from the original analysis, respectively). Interestingly, the LRE to re-assigned deme *189* (deme *213*) now mirrors the migration arrow to this deme from `TreeMix`. This could indicate that the original location of this sample was initially wrong, but once this was corrected, we see the 'true' signal of long-range genetic similarity. Since `TreeMix` does not use any spatial information, the results from that method was originally unbiased to this misreporting. We see a similar (albeit slightly weaker) signal for the re-assigned deme *585*, which now shows a source area geographically close to demes *690,822,719,626* and in the Hudson Bay, reflecting its position in the tree inferred by `TreeMix`.

Of the remaining 4 demes, all four LREs reflect directions inferred in `TreeMix`.

### S2 Figures

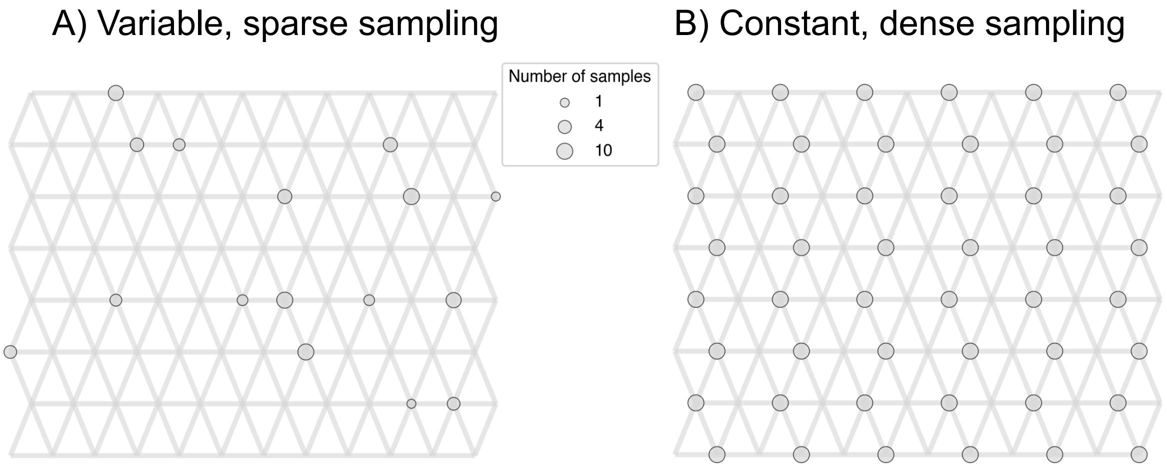


Figure S1: A visualization of the two sampling schemes from the main text for a single simulation replicate. The gray circles represent sampled demes on the overlaid triangular grid (as is custom in `FEEMS`, Marcus et al. 2021).

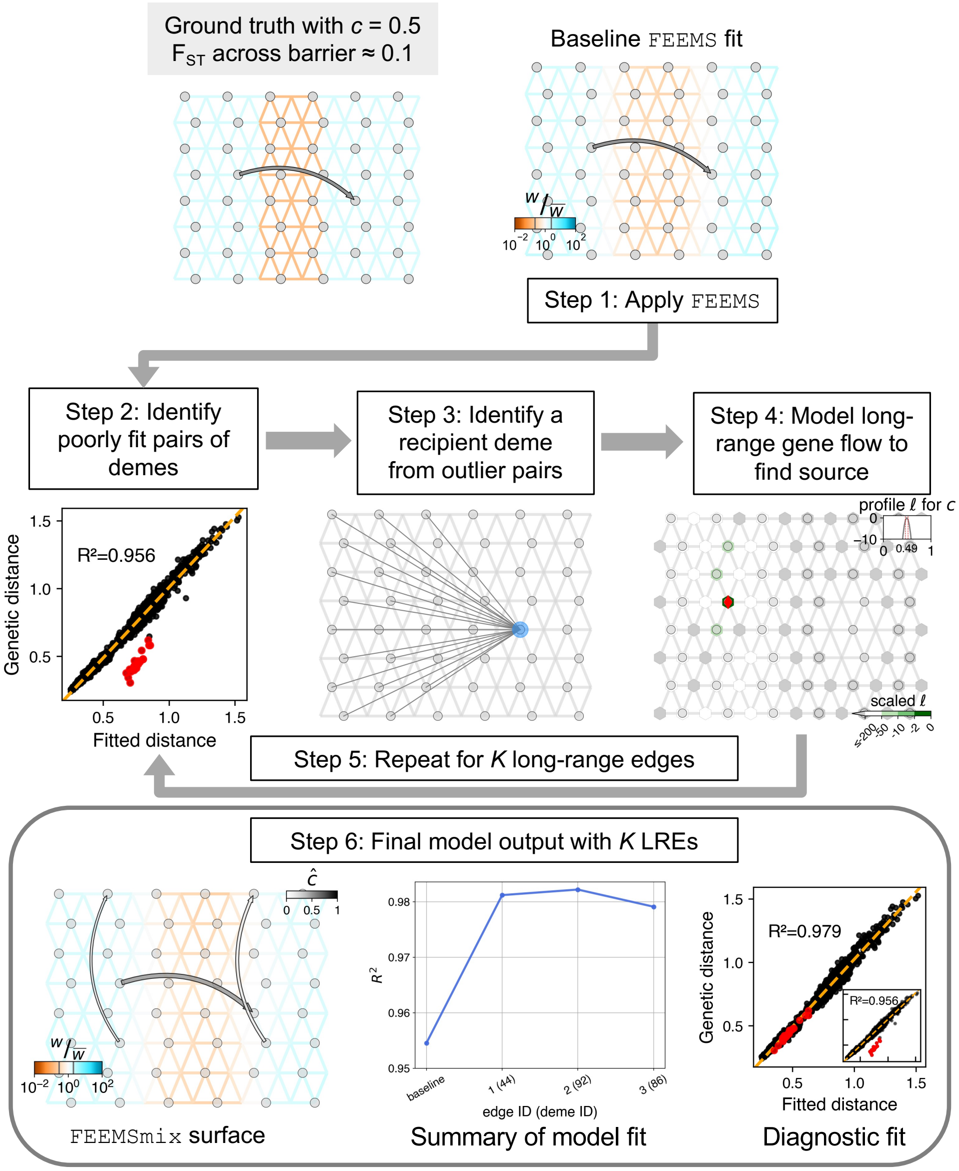


Figure S2: The *constant, dense sampling* analog to Figure 1 in the main text. With dense sampling, we see that the baseline **FEEMS** fit works well to capture the central barrier in the grid, with **FEEMSmix** also accurately estimating the source and strength of the long-range event. But we also see that adding too many edges can lead to overfitting with a decrease in model $R^2$ with $K = 3$ LREs.

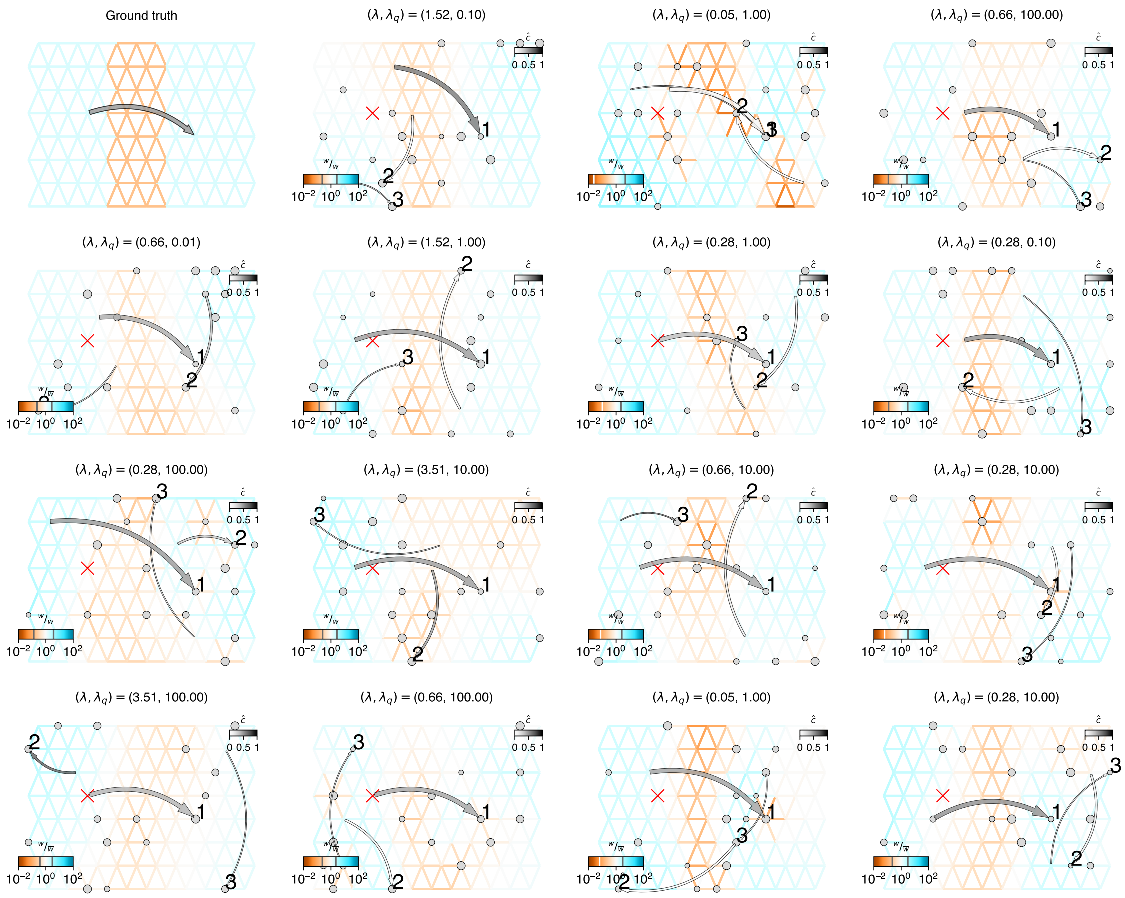


(A) Simulation replicates showing $K = 3$ LREs. The true source is marked as a red cross on the grid, and is never sampled as part of the strategy. Above each replicate is the value for the cross-validated $(\lambda, \lambda_q)$. The order the LREs are added in is indicated by the number at the destination deme.

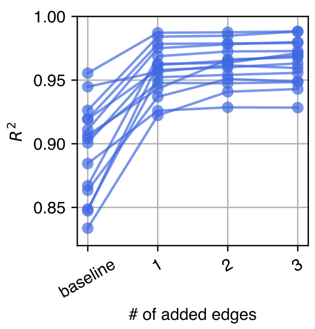


(B) Summary of model fits for each simulation replicate in subfigure A.

Figure S3: `FEEMSmix` results for 15 simulation replicates from the *variable, sparse sampling* scenario. In **A)**, we observe that the true destination deme is implicated as the first LRE in each replicate, while the other LREs are random with respect to the geographical source. In **B)**, we see a systematic increase in model $R^2$ with the first (true) LRE and a subsequent plateau with more added LREs.

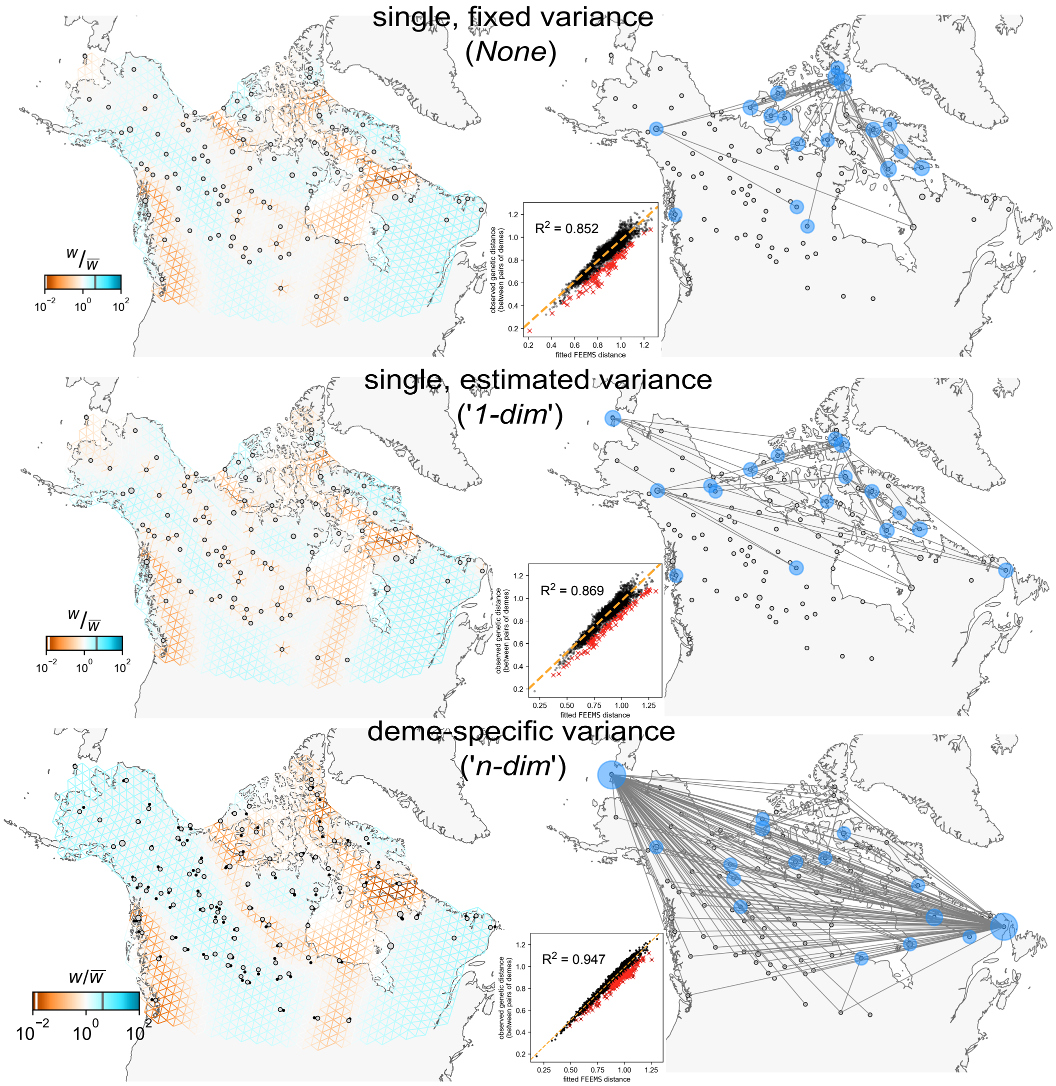


Figure S4: Baseline fits and residual outlier pairs from the three modes for how the variance parameters are modeled in `FEEMS`: *None* refers to a single variance parameter that is fixed at a value estimated by a model assuming a single weight across the entire grid (default in Marcus et al. 2021, comes from an initialization step before optimizing the weights, $\sim$ 2.1s), '*1-dim*' refers to a single, estimated variance parameter that is jointly estimated with all the other weights in the graph ($\sim$ 13.5s), and '*n-dim*' refers to estimating a variance parameter for each sampled deme (default in `FEEMSmix`, $\sim$ 11.3s). We observe that we obtain better fits with increasing number of parameters in the framework (based on $R^2$) though with increasing runtime. However, visually, all three methods pick up the major barriers and corridors in the data set. But, an important point to note here is how the pinwheel-like patterns around sampled demes disappear with the estimation of deme-specific variance parameters. Also, the outlier demes implicated in the fits change on a gradient between these modes: with Arctic and HighArctic demes being 75% of outliers in *None* to just 20% in '*n-dim*'.

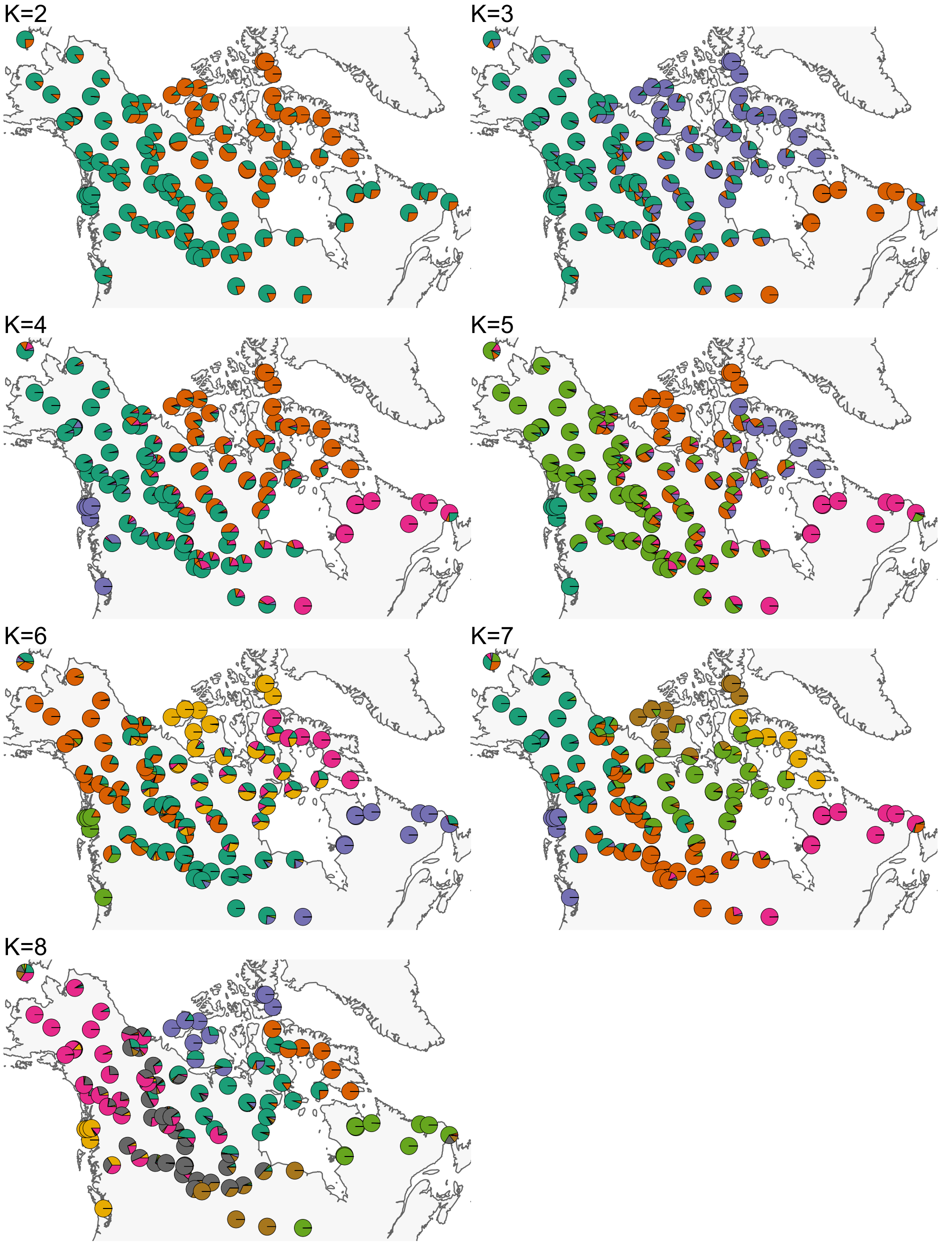


Figure S5: Average admixture fractions observed within each sampled deme from an `ADMIXTURE` (Alexander et al. 2009) analysis from $K = 2 - 8$.

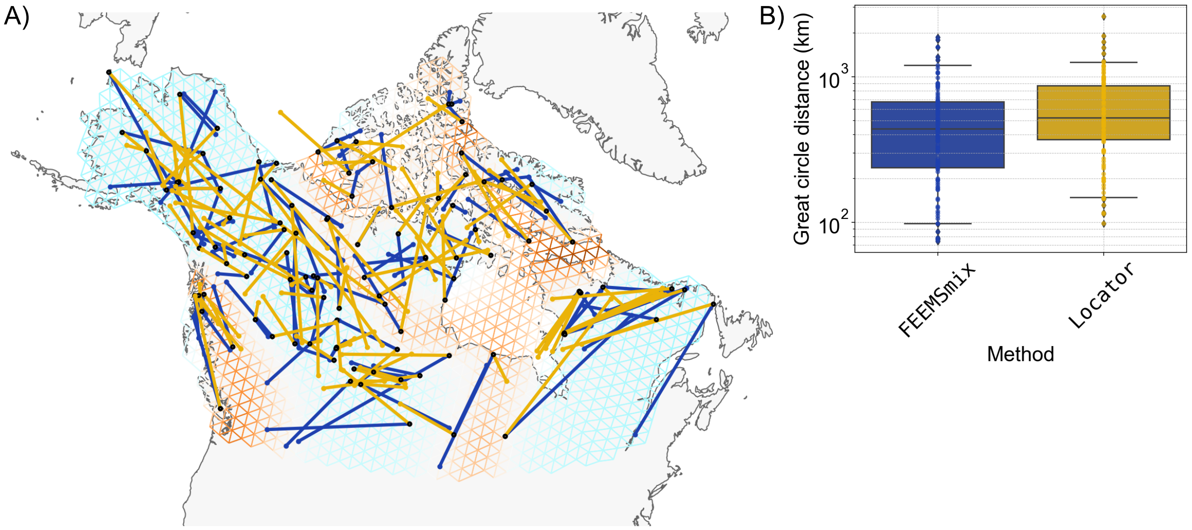


Figure S6: Comparison between `FEEMSmix` and `Locator` (a deep-learning based method from Battey et al. 2020) in predicting spatial locations of samples from the wolves data set in a leave-one-out based approach. We see comparable results between the two methods, with slightly better performance in `FEEMSmix` (decrease in median error of approx. 100 km, though high error with both methods indicate how mobile these wolves tend to be). True sample locations are shown as black points and the predicted locations are shown in the color corresponding to each method. An interesting point to note here is that the predicted locations from either method never cross the migration barriers as estimated by `FEEMS`.

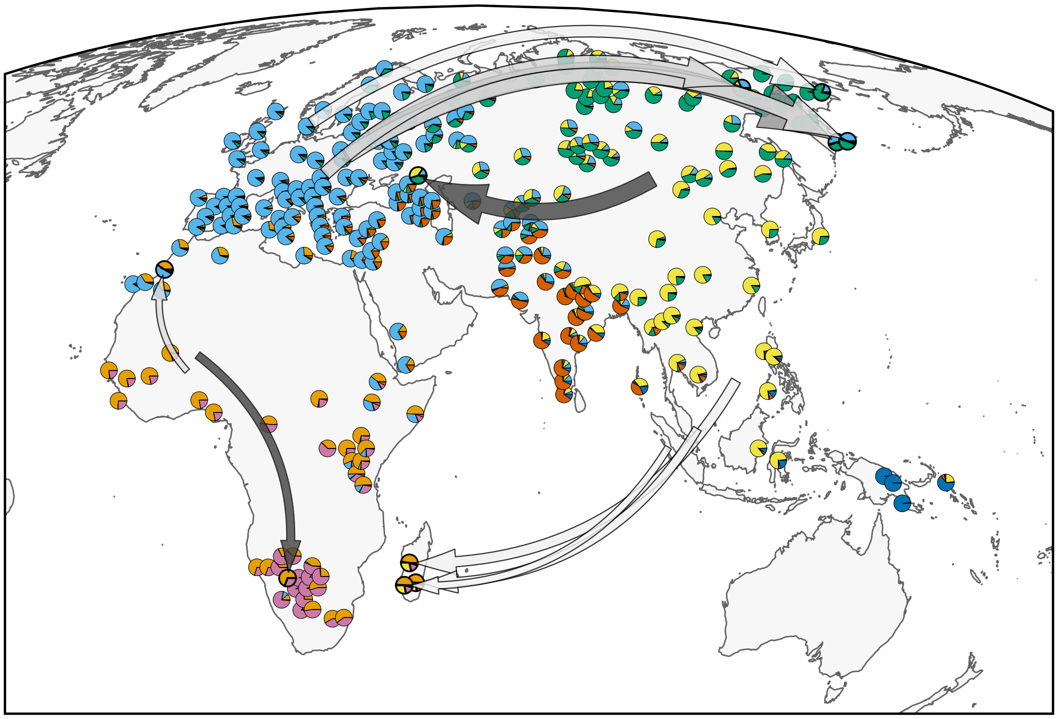

Figure S7: LREs from `FEEMSmix` overlaid on a map with `ADMIXTURE` results from a $K = 7$ analysis.

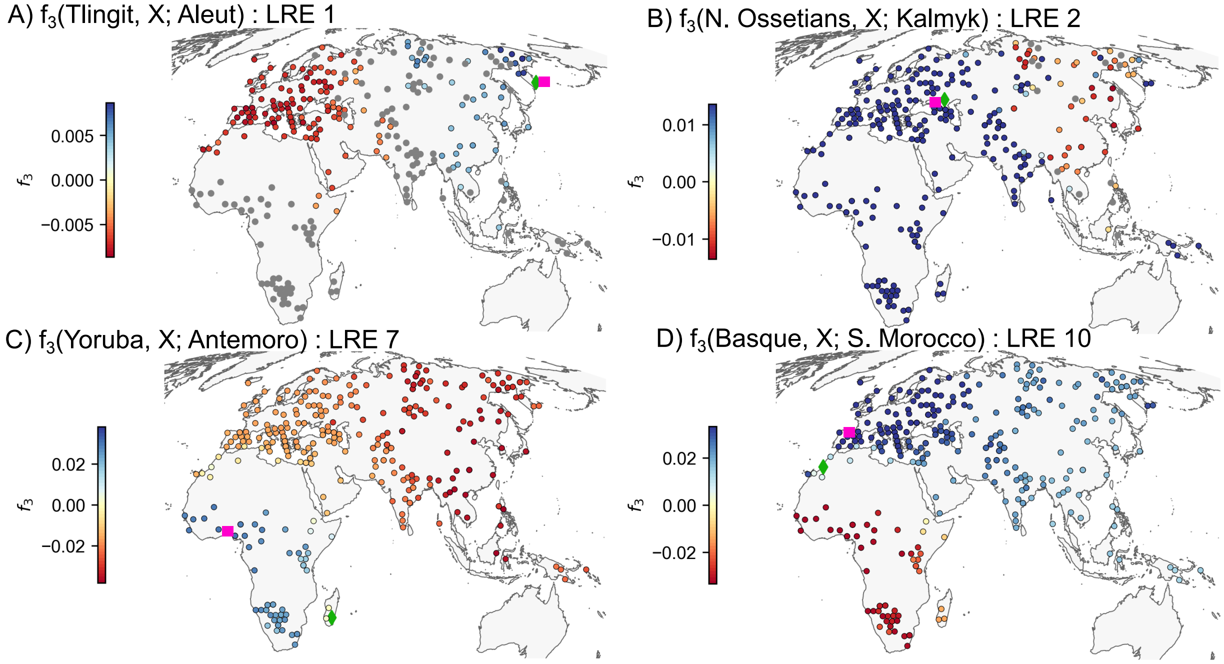


Figure S8: Admixture $f_3(A, X; T)$ statistics for selected putative destination demes identified by `FEEMSmix` (values computed using v651 of `AdmixTools` from Patterson et al. 2012). Here, $A$ is the reference population, $X$ is a testing population and $T$ is the target population (i.e., destination deme). We observe that populations with the most negative $f_3$ values (dark red) in each case are found in similar locations to the paired source demes inferred by `FEEMSmix` (in Figure 5B and C in the main text). The location of $T$ is displayed as a green diamond and the location of the corresponding $A$ is shown as a pink square.

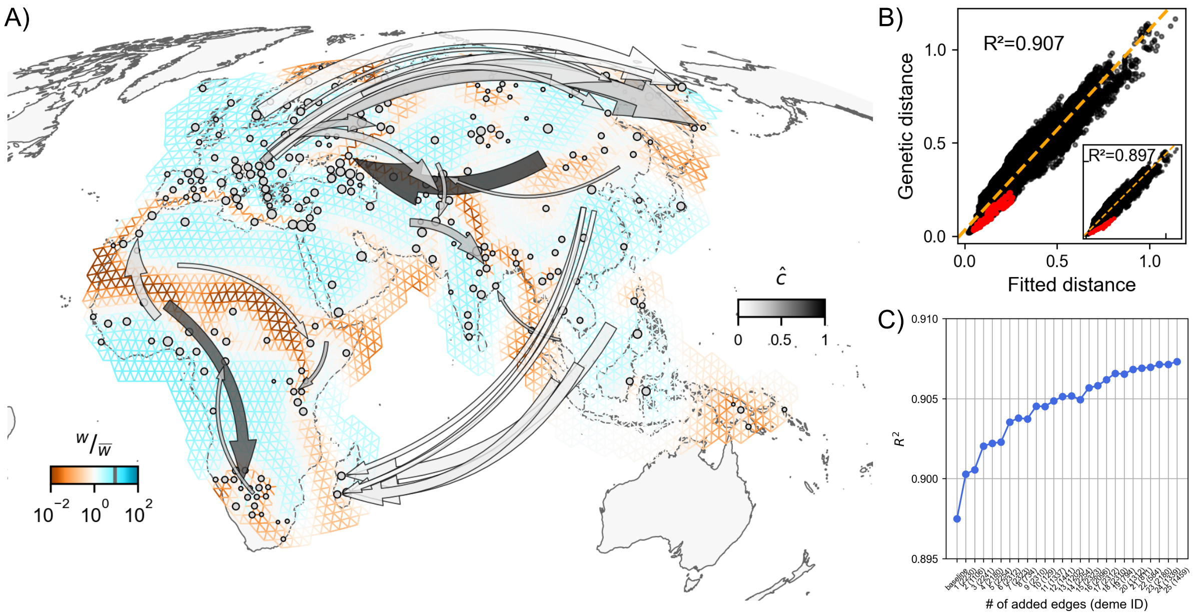


Figure S9: Full suite of `FEEMSmix` results from the human data set with $K = 25$ LREs.

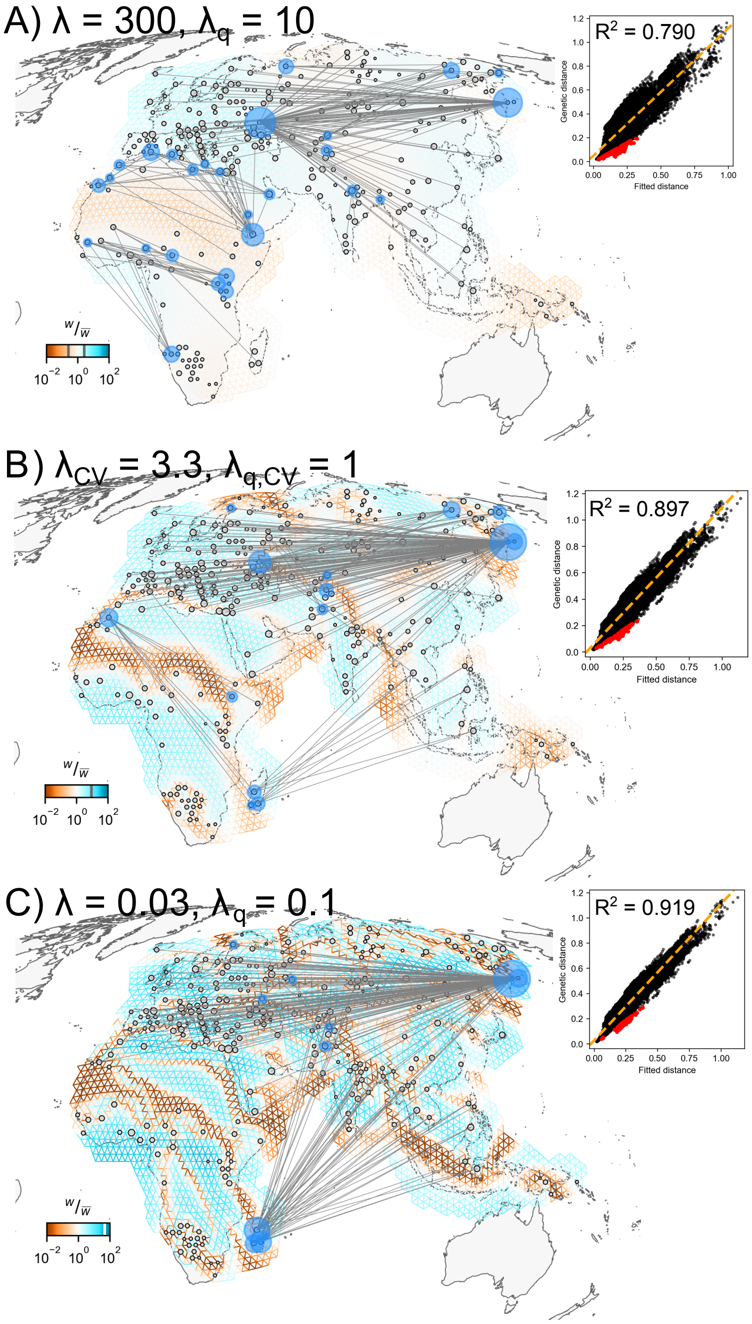


Figure S10: We present a range of fits to the human data set across different tuning parameter values, and show the top 1% of outliers using the method presented in the main text for human data. With decreasing $\lambda$ values, we see new demes being implicated as outliers in the fit, that disappear with a very low $\lambda$ value. This is likely due to overfitting with low $\lambda$, i.e., the baseline `FEEMS` fit connects two distant but similar demes via a snaking migration corridor connecting them, hence modeling away any residual. For instance, the Kalmyk individuals are no longer found as outliers, as they are connected to individuals further east through a migration corridor that punctuates a region of low inferred migration that is inferred with higher $\lambda$ values. As expected, the baseline model $R^2$ increases with decreasing $\lambda$, which is why leave-one-out cross validation is used to choose the optimal value for the $\lambda$'s.

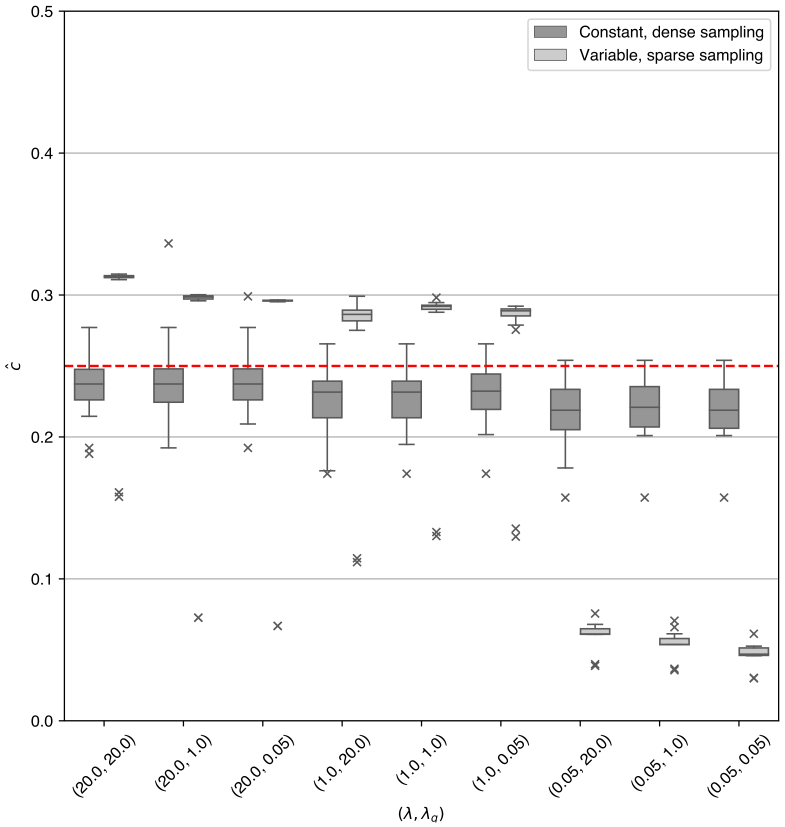


Figure S11: The effect of various tuning parameter values $(\lambda, \lambda_q)$ on the estimated admixture proportions $\hat{c}$ under the two sampling scenarios. The true simulated strength is marked as a red line at 0.25. We observe that there is little to no difference in estimated values across different tuning parameter combinations in the case of *constant, dense sampling*, but quite a big difference in the case of *variable, sparse sampling*. Essentially, for very small values of $\lambda$, FEEMS overfits the patterns in the data (see the more disjoint maps in Figure S3 for a visualization of this), leading to very little extra residual, and in turn, leading to an underestimation of the strength of long-range gene flow. The most common pair of tuning parameter values chosen by the cross-validation procedure was $(20.0, 100.0)$ for the dense scenario (similar to $(20.0, 20.0)$ in the figure) and $(0.3, 1.0)$ for the sparse scenario (similar to $(1.0, 1.0)$ in the figure).

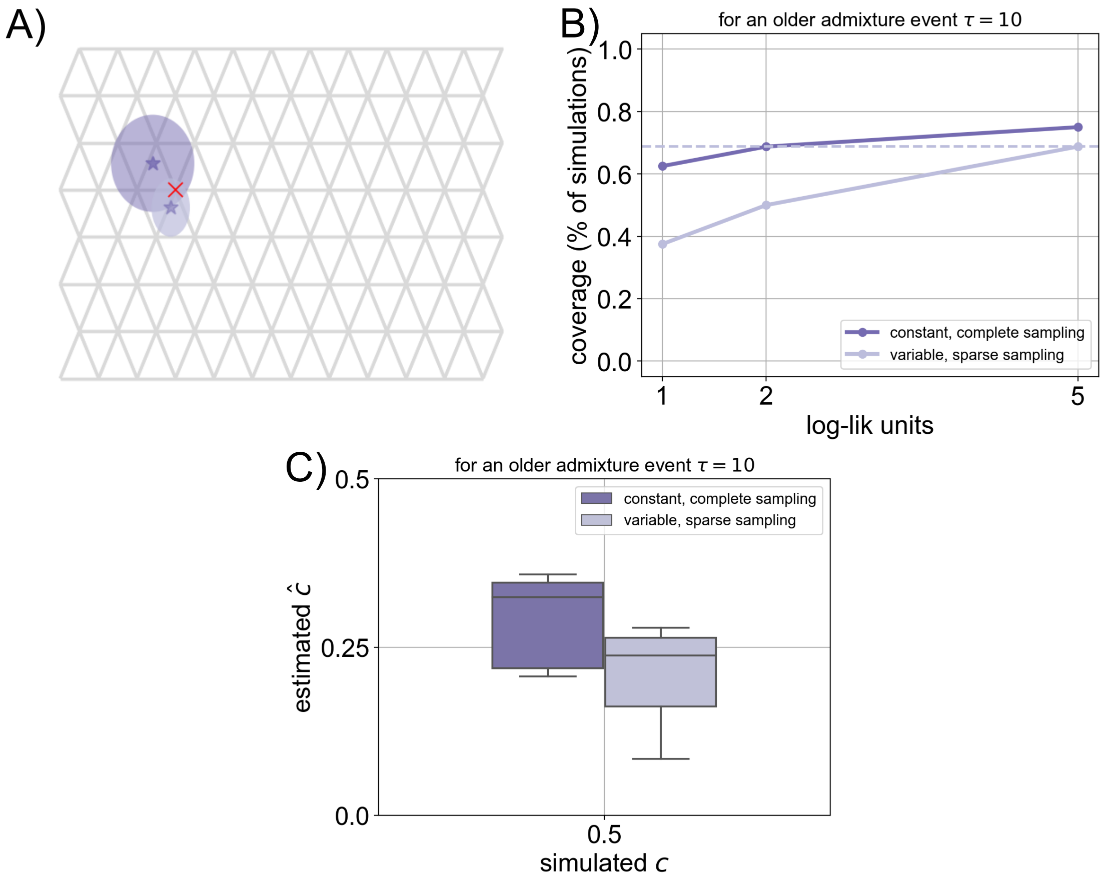


Figure S12: Result from a model mis-specification: inferring under a model of instantaneous admixture in the current generation (in `FEEMSmix`) for data simulated under an older admixture pulse 10 generations ago with $c = 0.5$. Any older admixture event will essentially diffuse the long-range migrant lineages across the landscape and leave a much less concentrated spatial signal. In **A)**, we see the mean inferred location (a $\star$) and $2\times$ standard errors of the mean (an ellipse) for the two sampling scenarios presented in the main text (compare to Figure 2A) . The true source in the simulations is represented by the red cross. In **B)**, we show the coverage statistics and see that the coverage statistics are lower in the case of an older admixture event (compare to Figure 2C). In **C)**, we see the model estimates a smaller admixture proportion than the one simulated since this signal is expected to decay over time due to the ongoing background migration on the landscape.

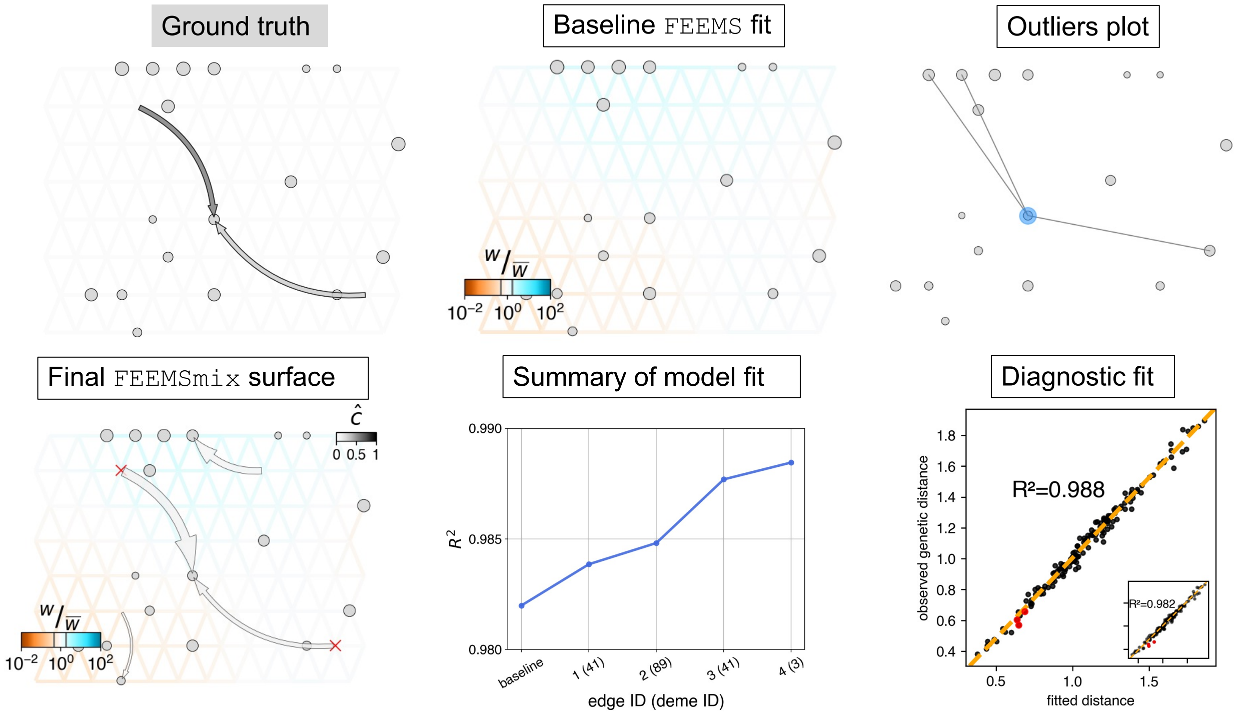


Figure S13: Result from a simulation with two geographically distant *unsampled* sources to the same destination over a uniform migration surface. In **Ground truth**, we show that there are two simulated long-range gene flow events with differing strengths (NW source: $c_1 = 0.5$, SE source: $c_2 = 0.25$) from diagonally opposite parts of the habitat. We see the two regions with the true sources (in red crosses) is found exactly by `FEEMSmix` on adding $K = 3$ LREs, though we also find that the true strength of the admixture is quite underestimated ($\hat{c}_1 \approx 0.1, \hat{c}_2 \approx 0.08$). This type of simulation also brings up the issue of power.

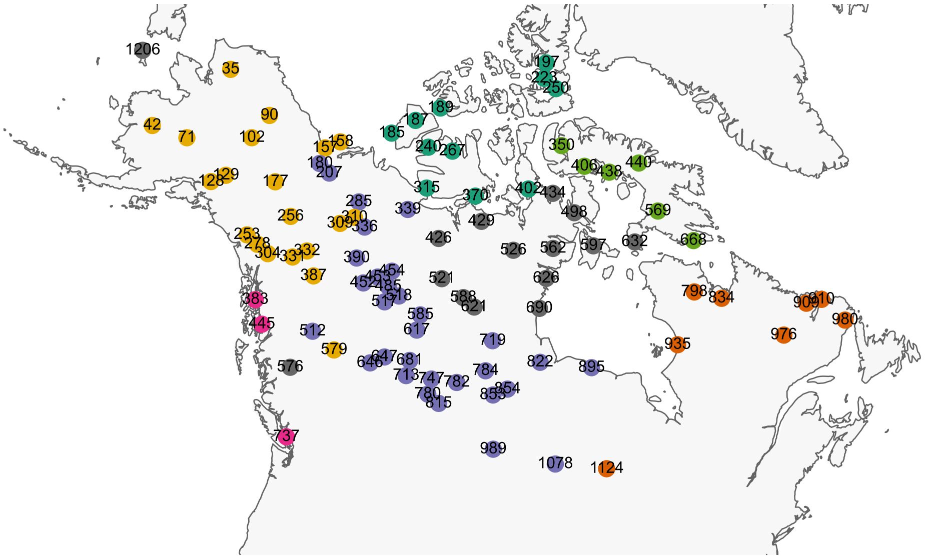

(A) A map of the wolves samples identified by the deme IDs used in the main text and colored by the 'Ecotype' classification from Schweizer et al. (2016).

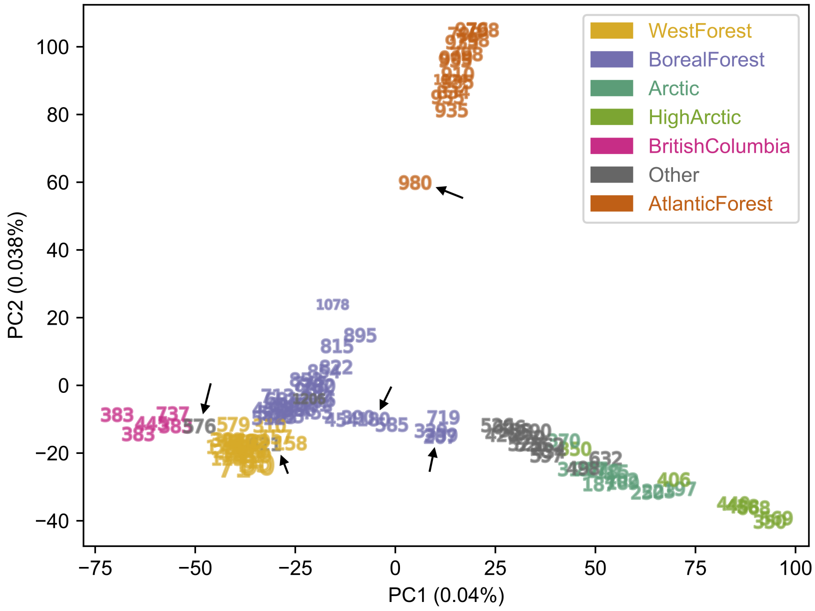


(B) A plot of the first two principal component axes on samples labeled by their deme IDs in `FEEMS` and colored by their 'Ecotype' classification.

Figure S14: Two plots to aid in interpretation of the `FEEMSmix` and `SpaceMix` results, and in identifying demes that are picked as recipient demes by these two methods (marked by black arrows). Deme *576* (also marked with a black arrow) might visually appear to be an outlier here, but its placement in PC space is in accordance with its geographic position.

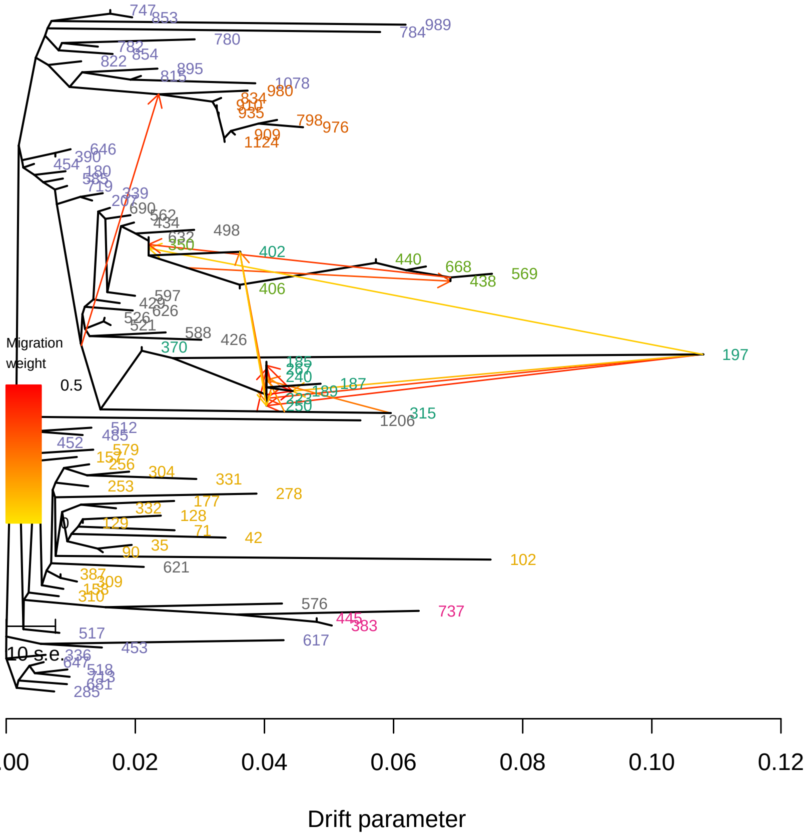


(A) Inferred topology with $m = 15$ migration edges

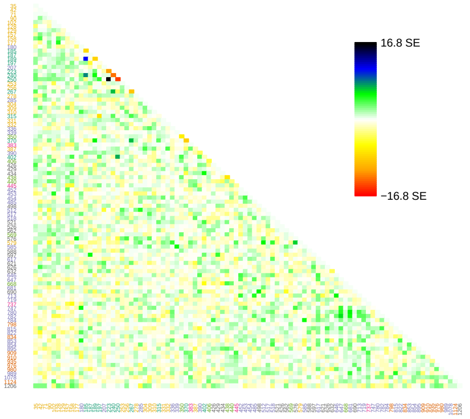


(B) Final residual matrix

Figure S15: `TreeMix` (Pickrell and Pritchard 2012) results with $m = 15$.

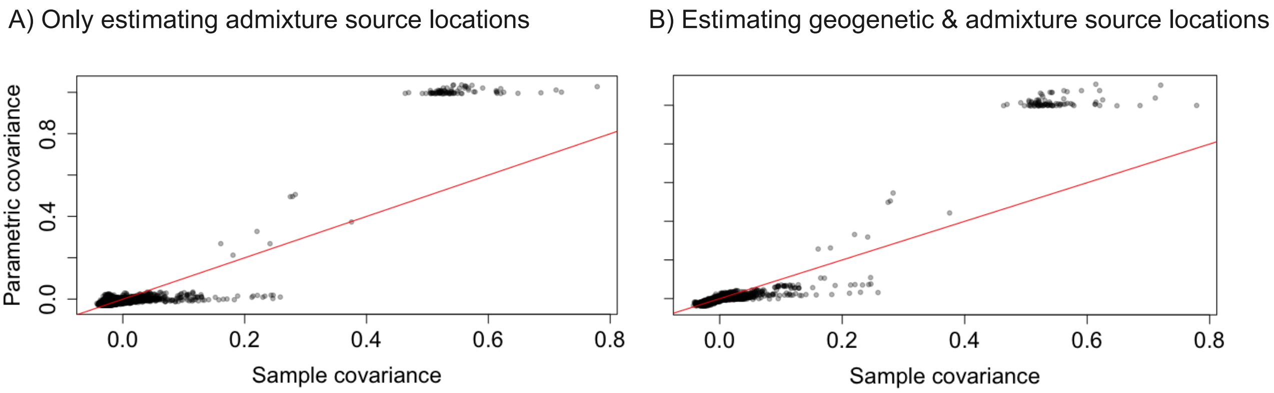


Figure S16: `SpaceMix` outputs showing pairwise sample (observed) and parametric (fitted) covariance across the two modes (A and B) of running the method mentioned in the text. Both modes produce $R^2 > 0.9$, and show a somewhat step-like pattern in the fits.

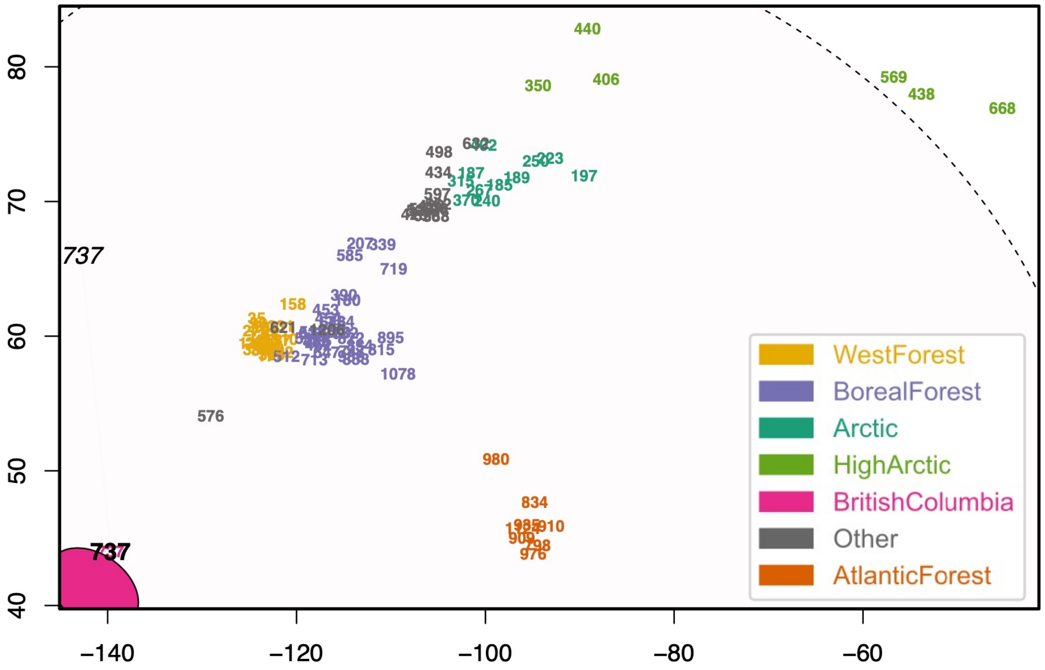


Figure S17: `SpaceMix` result when estimating both geogenetic and admixture source locations. We see that the two spatial axes of location places these demes in a manner similar to PC1 & PC2 in Figure S14B. The location of the sampled demes correlates with their 'Ecotypic' classification (in Figure S14A). Only deme *737* is implicated in an admixture event, with the location of the source extending over the entire habitat. In *all* cases, we see that the 95% credible intervals for the source locations span the entire 'geo-genetic' space with the maximum a posteriori estimates differing from the results from running a different mode in Figure S18.

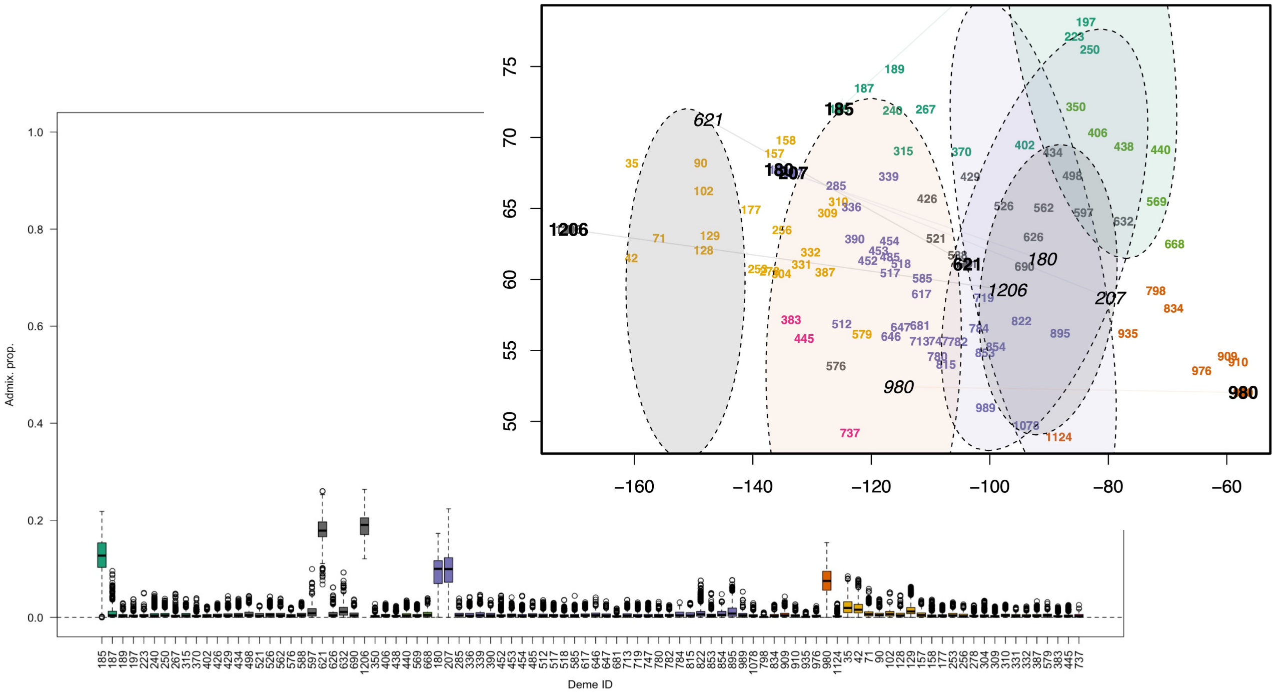


Figure S18: `SpaceMix` output when estimating only admixture source locations, keeping sampling coordinates fixed. In the inset plot, we see similar results with `FEEMSmix`, both in terms of the identity of the putative destination demes and the location of the sources of these admixture events. This similarity is also consistent with the results from `ADMIXTURE`. We also display the posterior distributions of the admixture proportions (as boxplots) of the events shown in ellipses in the inset plot.

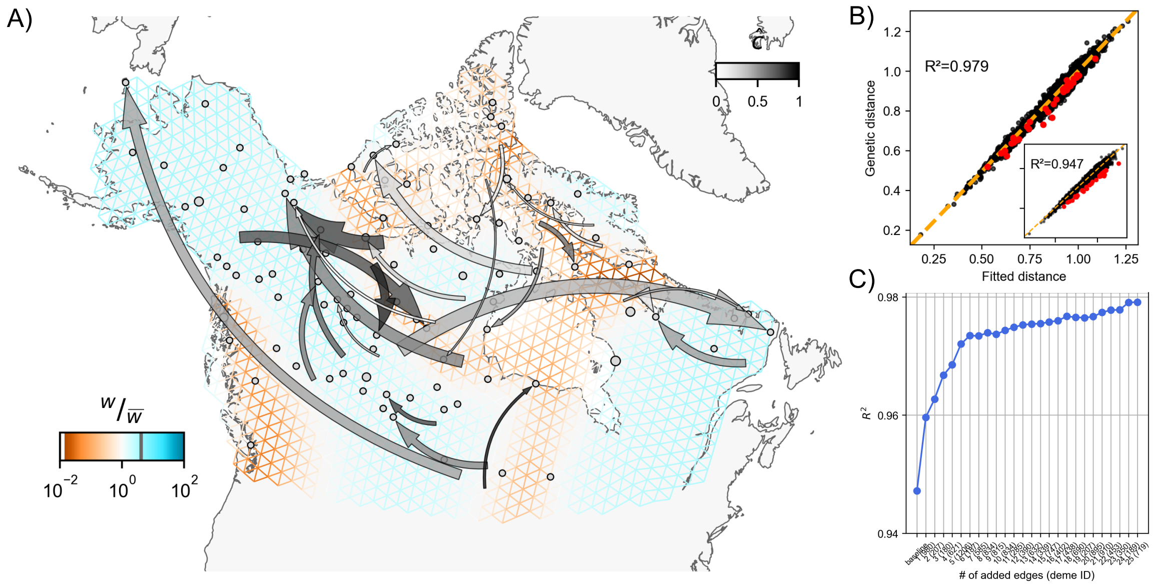


Figure S19: Full suite of FEEMSmix results from the wolves data set with $K = 25$ LREs.

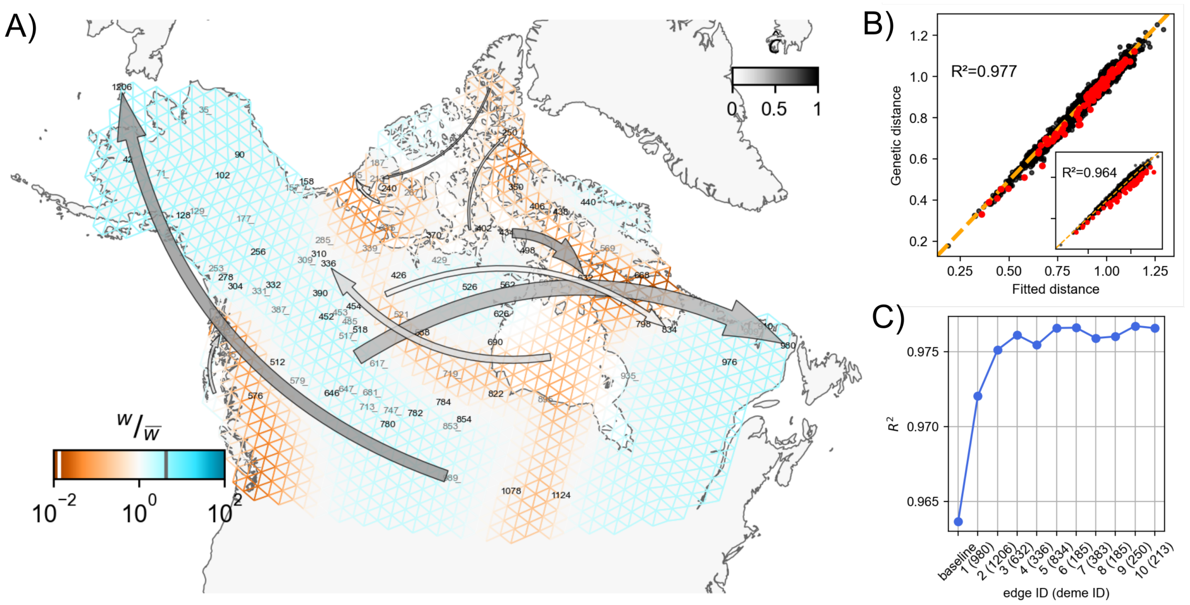


Figure S20: Full suite of FEEMSmix results with $K = 10$ from a (re)analysis of the wolf samples on correcting the misreported locations and removing the ambiguous samples.
